# Tuning the potency and therapeutic window of ImmTAC molecules by affinity modulation

**DOI:** 10.1101/2022.11.03.511676

**Authors:** Ian B Robertson, Rachel Mulvaney, Nele Dieckmann, Alessio Vantellini, Martina Canestraro, Francesca Amicarella, Ronan O’Dwyer, David K. Cole, Stephen Harper, Omer Dushek, Peter Kirk

## Abstract

T cell engaging bispecifics have great clinical potential for the treatment of cancer and infectious diseases. The binding affinity and kinetics of a bispecific molecule for both target and T cell CD3 have substantial effects on potency and specificity, but the rules governing these relationships are not fully understood. Using ImmTAC (Immune mobilizing monoclonal TCRs Against Cancer) molecules as a model, we explored the impact of altering affinity for target and CD3 on the potency and specificity of the re-directed T cell response. This class of bispecifics, exemplified by tebentafusp which has recently shown survival benefit in a randomized phase 3 clinical trial^1^, bind specific target peptides presented by human leukocyte antigen (HLA) on the cell surface *via* an affinity-enhanced T cell receptor and can redirect T cell activation with an anti-CD3 effector moiety. The data reveal that combining a strong affinity TCR with an intermediate affinity anti-CD3 results in optimal T cell activation, while strong affinity of both targeting and effector domains significantly reduces efficacy. Moreover, by optimising the affinity of both parts of the molecule, it is possible to improve the therapeutic window. These results could be effectively modelled based on kinetic proof-reading with limited signalling. This model explained the experimental observation that strong binding at both ends of the molecules leads to reduced activity, through very stable target-bispecific-effector complexes leading to CD3 entering a non-signalling dark-state. These findings have important implications for the design of anti-CD3 based bispecifics with optimal biophysical parameters for both activity and specificity.

**Graphical abstract:** 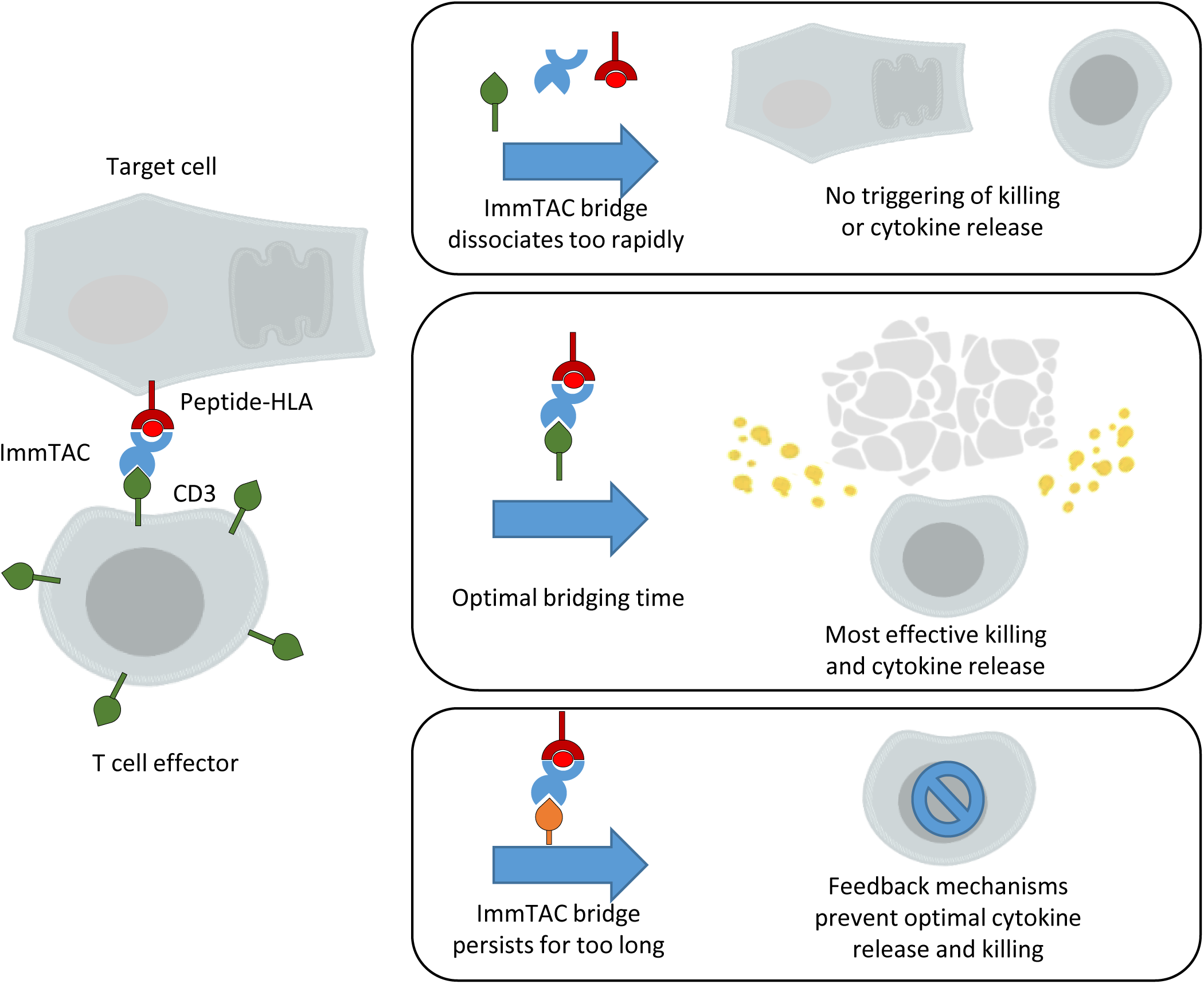

## Introduction

T cells are highly potent components of the adaptive immune system that, because of their ability to directly kill aberrant cells, have been recently targeted in several immunotherapy approaches for both cancer and infectious diseases. Target recognition by T cells is governed by the clonally expressed T cell receptor (TCR), which can initiate T cell activation upon binding to a cognate peptide-human leukocyte antigen (pHLA) complex, leading to immune synapse formation, phosphorylation of the CD3 signalling complex and downstream signalling.^2–5^ Importantly, T cells are highly sensitive, and can be activated by very low numbers of cognate pHLA (in the 10s per cell).^4, 6, 7^ Despite their exquisite sensitivity for antigen, TCRs bind to pHLA complex with weak affinity (K_D_s in the low µM range) and fast kinetics (half-life in seconds).^8^ T cells discriminate between different pHLAs on the basis of the duration of their interaction with the TCR,^9^ a property partly explained by the kinetic proofreading model,^10^ whereby the complex of pHLA and TCR must endure for long enough to initiate productive signalling.

ImmTAC (Immune mobilizing monoclonal TCRs Against Cancer) molecules have been developed utilising affinity-enhanced TCRs with high specificity towards target pHLA antigens. By forming bridging interactions between pHLA on target cells, and the CD3 signalling complex on the surface of T cells, ImmTAC molecules can drive formation of an immune synapse^11, 12^ which can mimic the ability of T cells to recognise pHLA (Figure 1A).^13^ The affinity of the anti-CD3 effector moiety is among the parameters that are likely to have a substantial effect on T cell redirection by bispecific T cell engagers, but there have been few studies systematically characterizing this effect across a broad range of affinities, or relating it to current models of T cell activation. Several studies have demonstrated that T cells respond best when they are stimulated by receptors or bispecific molecules with certain optimal affinities and kinetics, often preferring fast on-rate interactions over those with slow dissociation rates^14^. The detailed mechanisms behind this preference have not been completely elucidated, but several negative feedback mechanisms, such as phosphorylation by inhibitory Src family kinases^1^ and recruitment of specific phosphatases^3^, are candidates for converting the TCR-CD3 complex to a signalling-impotent dark-state after a prolonged period of activation^15^.

**Figure 1.**
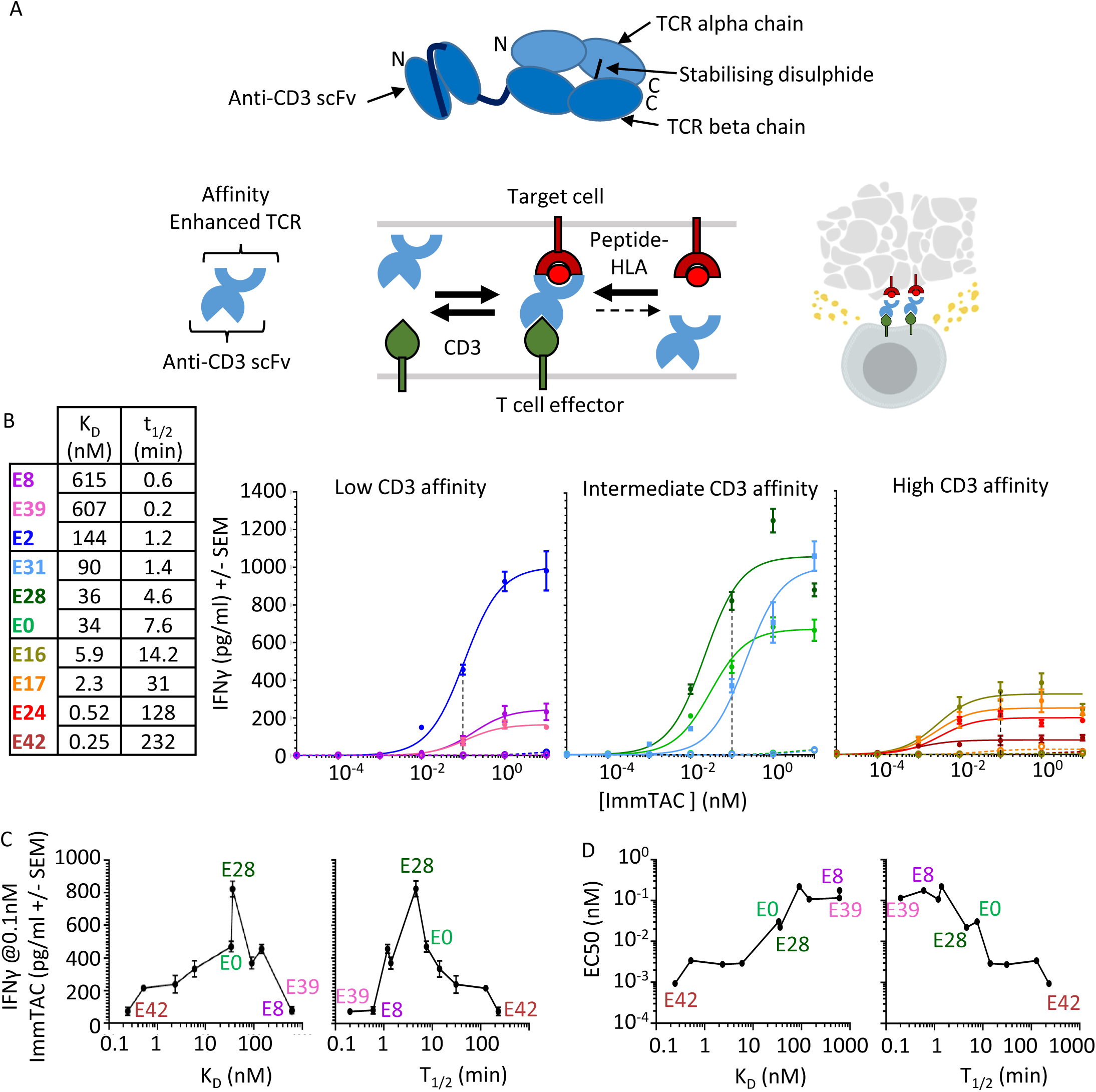
Effect of CD3 affinity on ImmTAC activity with a highly specific TCR (NY-BR-1). **A)** Summary of the basic structure of an ImmTAC molecule (top) and the general mechanism of action (below): An affinity matured TCR linked to an anti-CD3 scFv, forms bridges between effector T cells and target cells through strong binding to peptide-HLA with slow dissociation rates and intermediate binding affinity to CD3. This allows activation and redirection of T cells to specifically kill target cells. **B)** IFNγ ELISA of response of HLA-A*02:01 negative PBMCs in the presence of ImmTAC molecules and T2 cells pulsed with 5 nM Target peptide (solid lines and full circles), or without pulsed peptide (dotted lines and open circles). Three plots are shown for low, medium and high affinity anti-CD3 variants. Data points represent the mean of 8 different wells measured from a 384 well format plate. Black dashed line highlights the 0.1nM ImmTAC concentration used for summary plots of ELISA response efficacy shown in **C)** while potency (EC50) is shown in **D).** Both are plotted against CD3 affinity (left) or t1/2 (right) of CD3 binding.

Here, we used ImmTAC molecules as a model system to investigate the effect of modulating affinity for both target and CD3 on the potency and specificity of the redirected T cell response. By modifying the peptide ligand presented by HLA, we were able to use the same ImmTAC molecule to test a range of TCR affinities from µM to pM. Generation of a similar affinity range for the anti-CD3 effector moiety was achieved through targeted mutations in the paratope. Thus, this is the first study where a comprehensive landscape of affinity for both target (in this case pHLA) and CD3 has been explored, with focused characterisation of *in vitro* activity.

We then used bespoke mathematical modelling, based on previously established models incorporating kinetic proofreading^16^, to better understand the mechanisms governing our observations and to provide insights into how bispecific T cell engagers can be further optimised. Together, these data demonstrate the importance of a CD3 ‘dark state’ formed by the prolonged activation of CD3, which has major implications for selecting affinity combinations that maximise both potency and specificity. Modelling the kinetic mechanisms contributing to T cell activation also highlights bispecific combinations that might improve the therapeutic window, allowing the use of higher drug dosages with less risk of mimetic-driven cross-reactivity.

## Results

### Generation of anti-CD3 affinity variants

An existing scFv antibody^17^ that binds CD3 (E0 : K_D_ = 33.9 nM) was used as the basis for the generation of a panel of variants with a diverse range of binding affinities for CD3 (Table S1 and Figure S1 A). A site-directed alanine mutagenesis campaign guided by PDB 1XIW^18^ was used to produce a range of mutants with weaker affinity, while affinity maturation via phage display was employed to produce variants with stronger affinities (K_D_ = 615 nM (E8) to 0.25 nM (E42)). Additional site-directed mutagenesis combining both affinity enhancing and impairing mutations was used to further diversify the binding kinetics of the available anti-CD3 panel.

### Intermediate CD3 binding generates optimal efficacy for on-target activation

We fused anti-CD3 scFv variants to an affinity-enhanced TCR specific for the NY-BR-1_1106-1114_ peptide-HLA-A*02:01 complex^19^. The affinity of this TCR-pHLA interaction was K_D_ = 370 pM at 37°C as measured by SPR (Table S2 and Figure S1 B).

The activity of this ImmTAC panel was assessed in T cell redirection assays using PBMC from HLA-A*02:01 negative donors cultured with TAP-deficient T2 cells pulsed with target peptide, with IFNγ used as a readout of T cell activation. Very little cross-reactivity was observed when the ImmTAC molecules were titrated onto unpulsed T2 cells, even with the highest affinity anti-CD3 variants, demonstrating the absence of any relevant mimetic pHLA complexes in this system (Figure 1B and Figure S2).

The T cell redirection activity of bispecific molecules has two components; the potency, which represents the concentration of bispecific molecule required to observe activity, and the efficacy, which denotes the maximum level of activity observed. In the data presented here, potency and efficacy were not directly correlated: Variants with the weakest CD3 binding (E8 and E39) had low efficacy and potency, while the intermediate affinity variants, particularly E28, had the highest efficacy and good potency. The strong binding variants (E16, E17, E24, and E42), on the other hand, had the best potency but very poor efficacy.

Optimal efficacy can be seen with E2, E31, and E28 ImmTAC molecules, which have CD3 off-rates ranging from 1.2 – 4 mins and affinities of 144 –36 nM. E28 and E0 present a particularly interesting comparison as both had very similar affinities for CD3 (36 nM and 34 nM), but E28 had a faster on-rate and off-rate (t½ of 4.6 min *versus* 7.6 min for E0) and consistently gave a higher maximum response (Figure 1C-D). E8 and E39 were also similar in overall affinity (615 nM and 607 nM), but E8 had a slower off-rate (t½ of 0.6 min *versus* 0.2 min for E39) and outperformed E39, particularly in the more sensitive ELISpot assays (Figure S2). This suggests off-rate is a more useful predictor of activity than affinity alone.

### Optimal anti-CD3 affinity is dependent on TCR-pHLA affinity

The observed kinetic optimum for CD3 engagement suggested an optimal duration for the pHLA:ImmTAC:CD3 bridging interaction, consistent with the optimal dwell time observed for conventional pHLA:TCR interactions^15, 20, 21^. Therefore, we hypothesised that T cell activation by an ImmTAC molecule with high affinity for both pHLA and CD3 might be increased by weakening the binding to pHLA. To test this, a range of peptide mimetics of NY-BR-1 were identified that, when presented in the context of HLA-A*02:01, bound to the TCR with affinities ranging from K_D_ = 1.8 nM to 500 µM (Table S2 and Figure S1 B). These mimetic peptides were assessed for stability using the NetMHC 4 server^22^ and by confirming the presence of canonical anchor residues.

Four anti-CD3 variants with strong (E42), weak (E8), and intermediate (E0 and E28) binding to CD3 were tested on T2 cells pulsed with the NY-BR1 peptide and the mimetic peptides. Consistent with our hypothesis, weakening the affinity of the TCR-pHLA interaction increased the efficacy of E42 ImmTAC, while reducing both the efficacy and potency of E0, E28 and E8 ImmTAC molecules (Figure 2). When TCR affinity was weakened to K_D_ = 87 nM (MimC peptide), E42 significantly outperformed the other variants. This remained true for all weaker TCR affinities up to 1-2 _μ_M (MimF and MimG), where activity was very low, even with E42. No activity was observed with the MimH mimetic (K_D_ = 66 _μ_M) or MimI (K_D_ = 567 _μ_M; data not shown) with any anti-CD3 variant, suggesting an activity cut-off in the low micromolar range in this instance.

**Figure 2.**
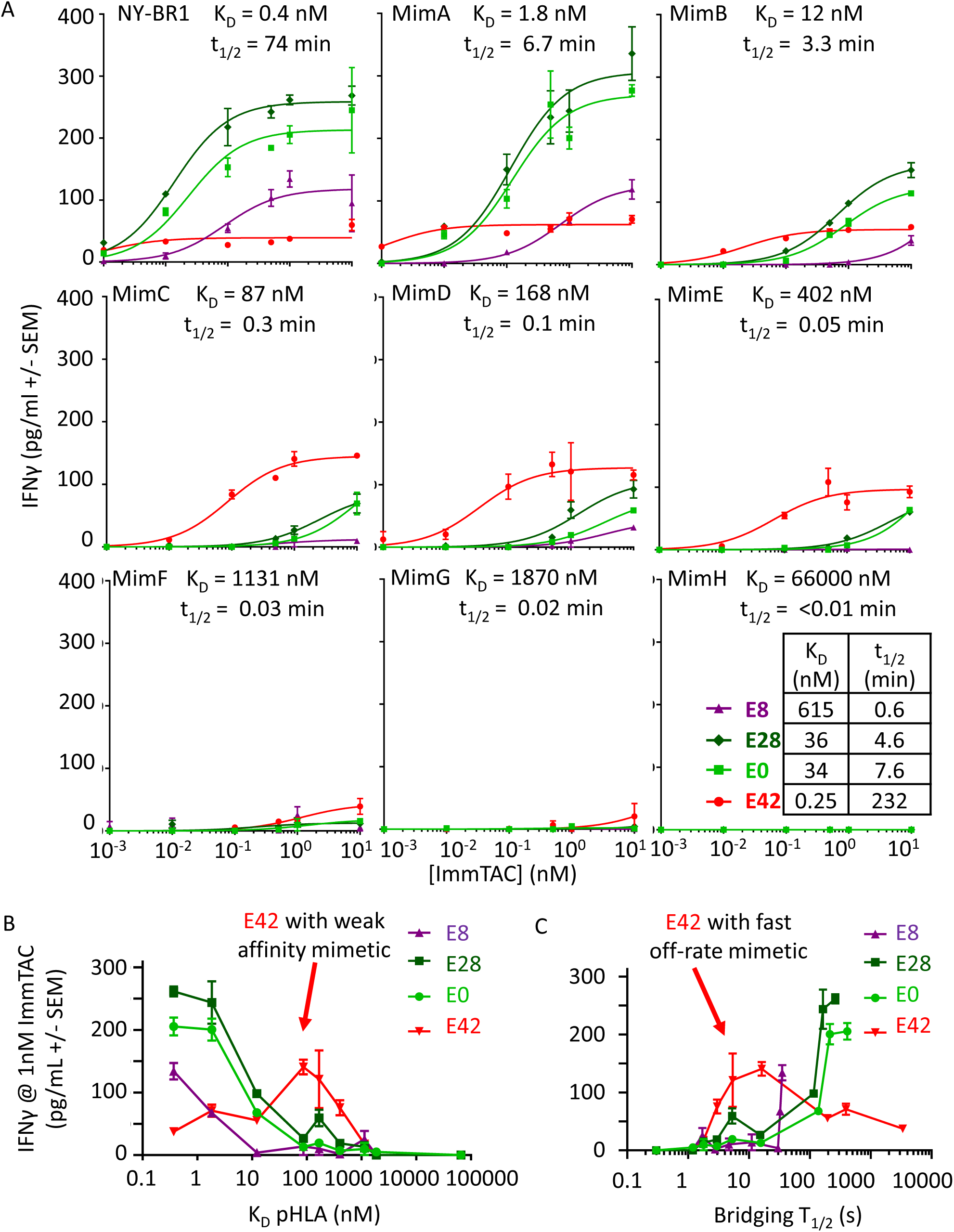
Effect of different combinations of TCR affinity and anti-CD3 affinity: **A)** NY-BR-1 target peptide and a range of its mimetics were added to 50,000 T2 cells at a concentration of 5 nM, (37 ^0^C affinities of each peptide HLA for TCR shown in the figure). Four ImmTAC molecules with E8, E28, E0, and E42 anti CD3 arms, were added at varying concentrations along with 40,000 PBMC cells. IFNγ was then measured by ELISA after a 48 hr incubation in a 96 well format. **B)** Summary plot of ELISA data showing response to 1 nM ImmTAC plotted against TCR binding affinity to the different mimetic peptides. **C)** Summary plot of ELISA data showing response to 1 nM ImmTAC plotted against the combined dissociation rate of TCR and anti-CD3 (i.e. the estimated T½ of a cell-cell bridge).

Given that an ImmTAC bridge can break by unbinding from either pMHC or CD3, we plotted the ImmTAC efficacy over the bridge lifetime (1 / (pMHC koff + CD3 koff)) (Figure 2C). While the data for E42 indicates an optimal duration for the pHLA:ImmTAC:CD3 bridging interaction, E8, E0 and E28 show a different optimum, indicating that a simple assessment of the duration of the bridging complex alone is not sufficient to fully explain the data.

Similar effects were also observed with a commercial CD4+ Jurkat NFAT luciferase assay when these cells were incubated with the same ImmTACs and peptide pulsed T2 cells (Figure S2 D), confirming the effect is not specific to IFNγ release.

### Kinetic proofreading with limited signalling model fits observed T cell activation

To test whether a conventional model of T cell activation could help explain the experimental observations, a mathematical model was constructed based on previously published mechanisms.^15, 16^ Such a model can help simulate data out of reach of available experimental tools and further aid the design of highly optimised molecules depending on the targeted therapeutic niche.

Ten ordinary differential equations (ODEs) were defined based on an established ‘kinetic proof-reading with limited signalling’ model of T cell activation.^15^ This model invokes a non-signalling ‘dark state’ for CD3 when the bridging interaction is sustained beyond a certain duration. The system of equations takes the kinetic binding rates as inputs along with starting concentrations based on cell density, predicted epitope number, and ImmTAC concentration. Changes in local concentration of free receptors and receptor-ImmTAC complexes on the surface of T cells and target cells are then modelled accordingly (Figure 3). Several variables were incorporated into the model to account for unknown parameters such as the rate of kinetic proofreading and rate of dark state formation. All constant and variable inputs are summarised in Table S3.

**Figure 3.**
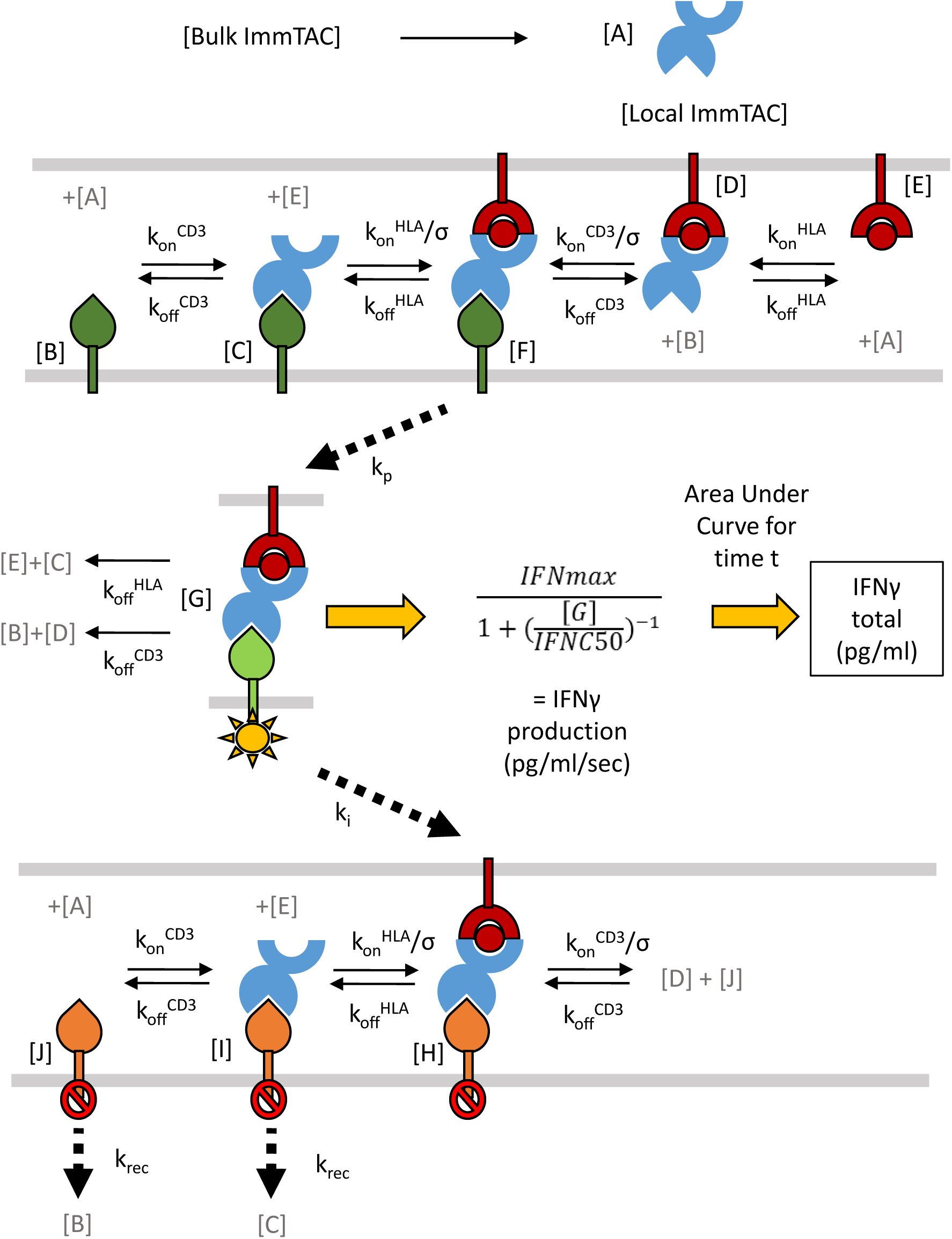
Structure of mathematical model for T cell activation. Ten local concentrations (A-J) were modelled using ten ordinary differential equations and these concentrations and their relationships are summarised here. Cell membranes are represented as grey lines, ImmTAC is shown in blue, peptide HLA in dark red, resting CD3 in dark green, activated CD3 in light green, and inhibited ‘dark state’ CD3 in orange. Relevant rate parameters are shown above reaction arrows and letters in grey denote involvement of other model components, (each component is only shown here once for simplicity). Concentration of active CD3 was translated into a rate of IFNγ production per second following a simple 3 parameter dose-response relationship, then the area under the curve fitted for total IFNγ production over the course of the experiment. The local free ImmTAC concentration (A) was replenished by rapid diffusion from an inexhaustible bulk solution.

To fit the model to the observed data, Approximate Bayesian Computation coupled with Sequential Monte Carlo (ABC-SMC) analysis was carried out on individual data sets. The model gave a good fit to the dataset presented in Figure 1B (Figure 4A), where only CD3 affinity was varied. For the dataset presented in Figure 2A, where affinity for both CD3 and for pHLA was varied, the model captured the key features of the data, with some deviation (Figure 4B). Probability distributions and ‘best particle’ parameters obtained for each experiment are shown in Figure S3 and Table S4. To test the robustness of these observations the output of the model was calculated after doubling individual constant or fitted parameters (Figure S4), demonstrating the pattern of behaviour was well conserved and would not be significantly altered by any errors in the measurement of binding kinetics.

**Figure 4.**
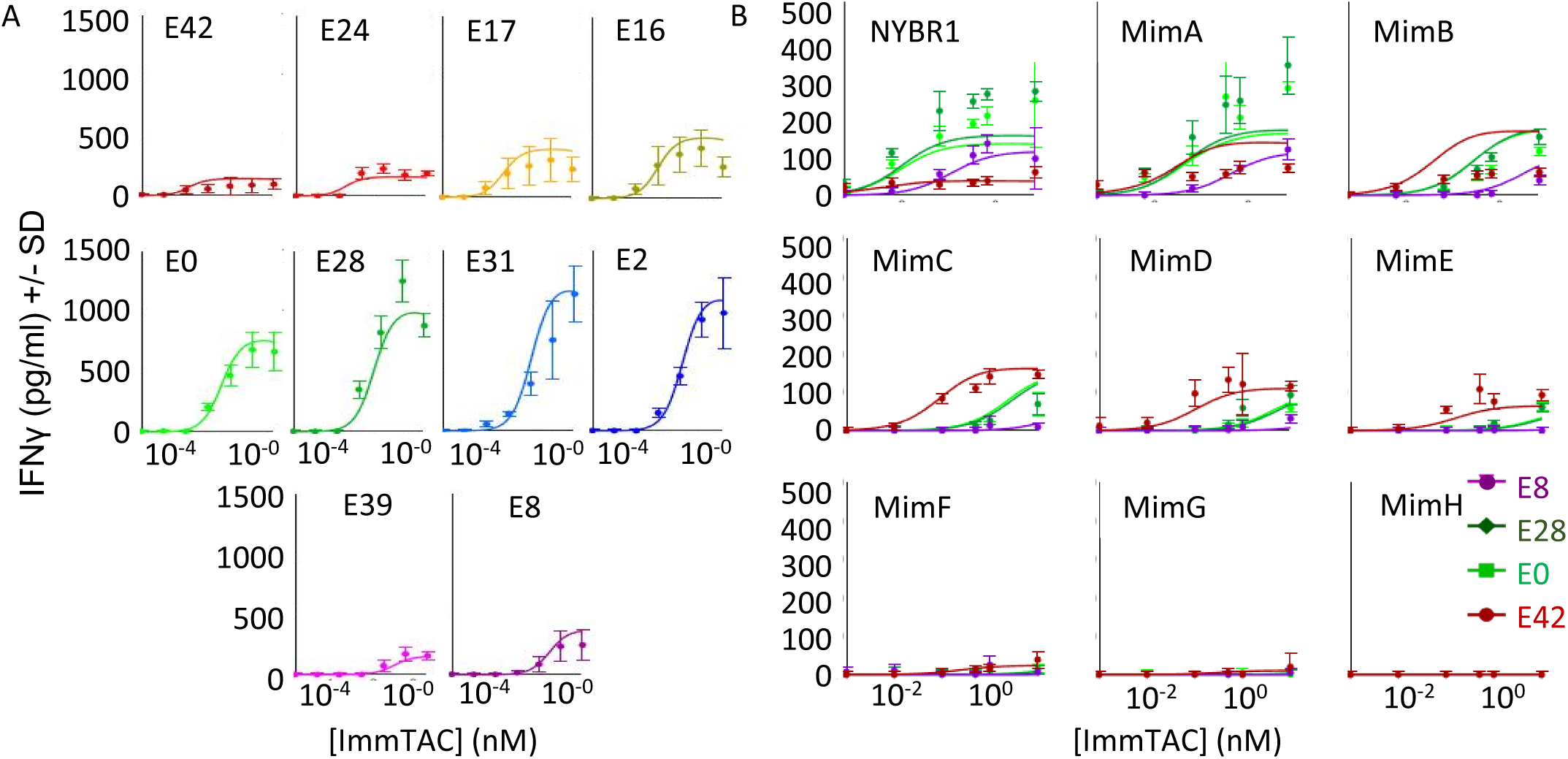
Model fitting results compared to the data. Fits of mathematical models applied to two data sets from Figures 1 and 2. Simulated data shown here as solid lines generated using ‘best particle’ values for final iteration of ABC-SMC runs (Table S4). **A)** Fit to ELISA data from Figure 1B generated with T2 cells pulsed with 5nM target peptide and 10 different anti-CD3 variants. Error bars represent standard deviation from 8 data points. Parameter values used in fit shown in Table S4 **B)** Fit to ELISA data from Figure 2 generated with four anti-CD3 variants on T2 cells pulsed nine different peptides at 5 nM. Error bars represent standard deviation over 3 data points. Parameter values used in fit shown in Supplementary Figure S2 and Table S4.

Using this model, T cell activation was simulated with a wide range of pHLA and CD3 binding kinetics. The parameter values fitted in Figure 4B (Table S4) were used as this represented the broadest dataset in terms of both CD3 and pHLA binding. When TCR affinity was strong (K_D_ = 100 pM), a clear optimum off-rate was seen for anti-CD3 variants (∼100s) past which strengthening CD3 binding significantly reduced efficacy with minimal improvements to potency (Figure 5A, left panel). However, with weak TCR binding (K_D_ = 1 µM), strengthening CD3 binding continuously improved potency without reducing efficacy (Figure 5A, right panel).

**Figure 5.**
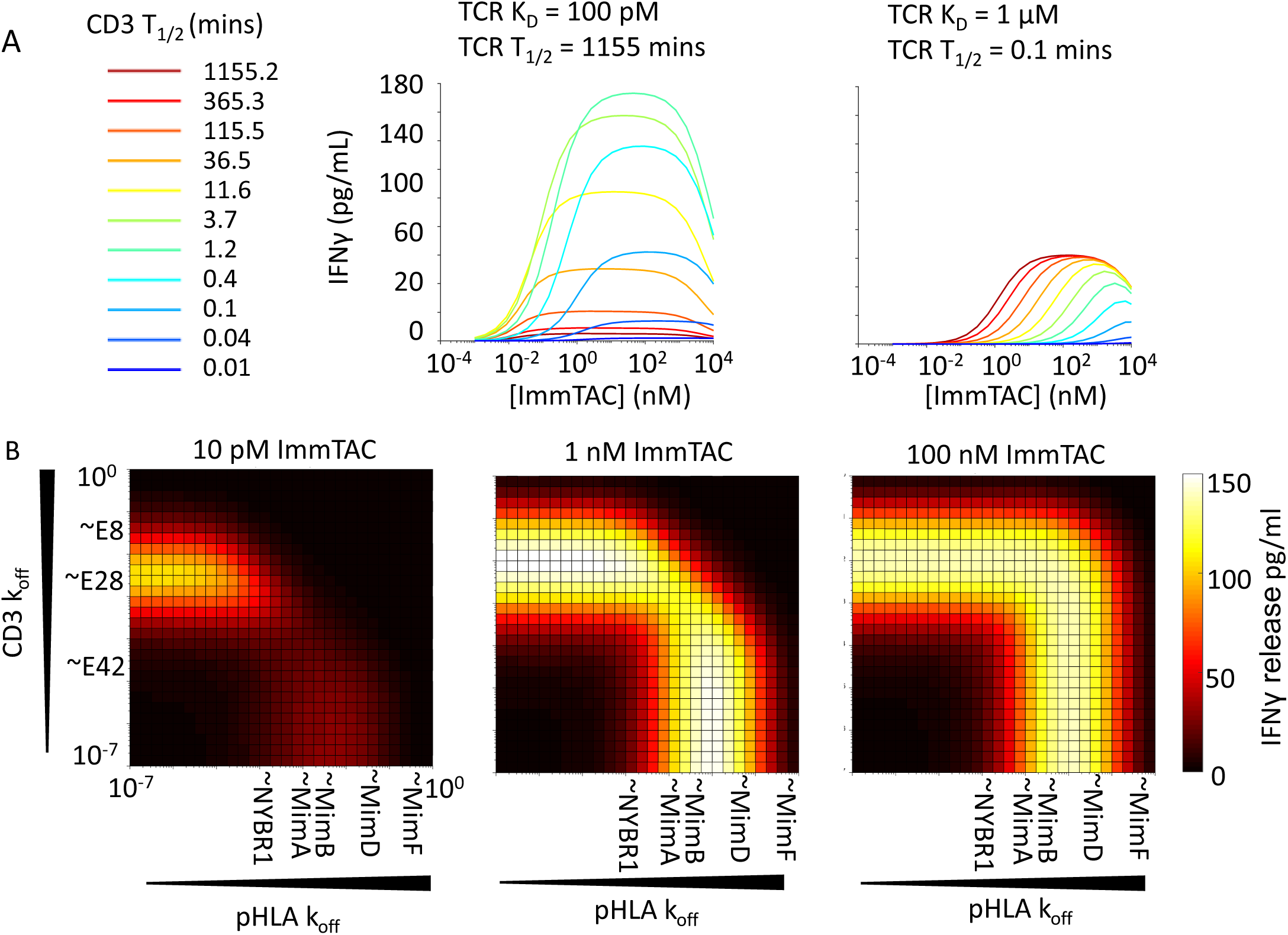
Implications of mathematical model for ImmTAC potency and efficacy. **A)** Effect of varying CD3 T_1/2_ with fixed k_on_ and long lived TCR-pHLA binding (T_1/2_ = 1155 mins) left, or short lived TCR-pHLA binding (T_1/2_ = 0.1 mins) right. **B)** Heat maps plotting predicted IFNγ responses with different CD3-HLA affinity combinations at various ImmTAC concentrations (as written above each heat map). Off-rates of anti-CD3 variants and TCR binding mimetics marked on axes for reference, although their on-rates also vary.

To better visualise the different effects of pHLA and CD3 off-rates on T cell responses, heat maps of predicted IFNγ release were constructed by varying the off-rate combinations employed at three different ImmTAC concentrations (Figure 5B). At low ImmTAC concentrations of 10 pM, combining strong pHLA binding with intermediate CD3 binding gave more response than combining strong CD3 binding with intermediate pHLA binding. This appears to be due to a better propensity for serial triggering with strong pHLA binding if, when dark state formation by both formats is equivalent, a bridging interaction is more likely to break to release ImmTAC bound to pHLA, which is then more primed to serially-trigger than an ImmTAC bound to a CD3 molecule still in the dark state. This benefit is lost if recovery from the dark state is made extremely fast (Figure S5), and above 1 nM ImmTAC strong pHLA binding had no clear advantage over strong CD3 binding. Combining both strong pHLA and strong CD3 binding gave no advantage at any concentration due to dark-state formation.

### Reducing CD3 affinity is predicted to widen the therapeutic window

As well as optimising potency, the different CD3 and pHLA binding optima demonstrated here provide an opportunity to select bispecifics with improved discrimination between high-affinity target and low-affinity mimetics. To model T cell responses when both on and off-target pHLA ligands are presented together, five ODEs were added to the mathematical model to describe local concentrations of free off-target pHLA, off-target ImmTAC complex, off-target-CD3 bridge, off-target active signalling, and off-target-dark-state bridge.

Simulations with this expanded model showed that when T cells were exposed to both a high affinity target peptide (K_D_ = 1 nM) and an intermediate affinity ‘mimetic’ (K_D_ = 100 nM), a hybrid response profile was obtained (Figure 6A). With strong CD3 binding (K_D_ = 0.1 nM – 10 nM), dark state formation suppresses on-target activity, limiting potency (Figure 6A, left), while at the same time potency on cells presenting the mimetic alone is increased (Figure 6A, right). This means that with a strong affinity TCR, strengthening CD3 affinity will shrink the therapeutic window if relevant mimetics are present, whereas weakening CD3 binding up to a point can widen this window.

**Figure 6.**
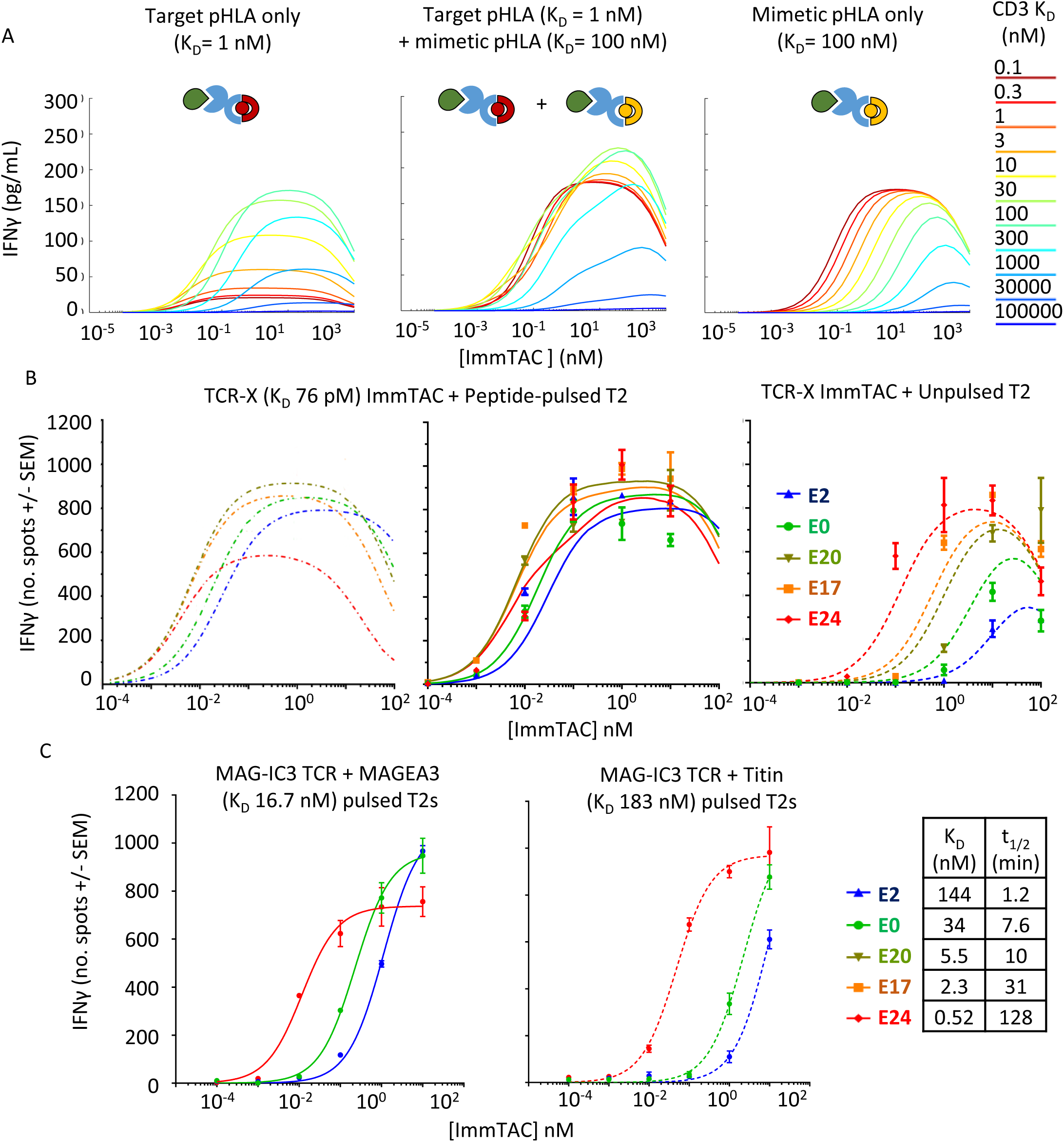
Effect of CD3 affinity on ImmTAC therapeutic window. **A)** Modelling of IFNγ response to two different HLA molecules (target and mimetic) was run with parameters fitted to NY-BR-1 data and an expanded 15 ODE model. Left: With only 1 nM target pHLA present, Middle: both 1 nM affinity target and 100 nM affinity mimetic pHLA molecules present, or right: only mimetic 100 nM affinity pHLA modelled. (The same presentation level of peptides used in both instances and on rate was kept constant at 0.1 μM^-1^ s^-1^). **B)** Modelling and data for IFNγ production from PBMCs in the presence of cross-reactive TCR-X ImmTAC and T2 cells pulsed with 5 nM Target peptide (left two graphs), or unpulsed cells (right). Lines show fits generated by the ODE model, without (left) or with (middle and right) fitted mimetic included in the model, and dots show ELISpot data points. Parameters of fit are shown in Figure S4. **C)** ELISpot data with MAG-IC3 ImmTAC and HLA-A1 transfected T2 cells pulsed with the MAGEA3 target peptide that binds the TCR with a K_D_ of 17 nM (left), or the mimetic Titin that binds the TCR with a K_D_ of 180 nM (right),(2.5 μM peptide pulse used in both instances). Curves represent 3-parameter fits. (Modelling was not performed for this TCR as data on peptide presentation level were not available)

To confirm whether the therapeutic window could also be increased based on the finding from the model, we collected data using an alternate affinity-enhanced TCR (TCR-X) that binds its target pHLA complex with a K_D_ of 76 pM at 37 °C (t½ = 325 min, Table S2). Unlike the NY-BR-1 specific TCR, TCR-X displays cross-reactivity towards unpulsed T2 cells and antigen-negative cells, allowing changes to the therapeutic window to be more readily examined.

ImmTAC molecules were made by fusing TCR-X with a select panel of anti-CD3 variants, then titrated onto TAP-deficient T2 cells that were either left unpulsed or pulsed with target peptide. T cell activation was assessed using an IFNγ ELISpot assay to maximise sensitivity (Figure 6B) and the new 15 ODE model was fitted to the experimental data (Figure S6 with fitted parameters in Table S5).

As predicted by the model, strengthening CD3 binding gave minimal improvement to on-target activity with this high affinity target, while cross-reactivity to unpulsed cells steadily increased (Fig 6B and S6), thus shrinking the therapeutic window. The effect was so striking that barely any window remained between target-pulsed and unpulsed cells with the strong CD3 binding E24 variant. The weaker binding E2 variant on the other hand gave some improvement to the therapeutic window relative to E0.

The identity of the mimetic peptide (or peptides) bound by TCR-X on unpulsed T2 cells is unknown, but as part of model fitting, a single ‘mimetic off-rate’ and ‘copies per cell’ were fitted to the data alongside other parameters. Using the data from one representative effector PBMC donor, the fitted parameters produced a mimetic off-rate of 0.015 s^-1^, (K_D_ = 30 nM), and 86 copies per cell (Table S5). However, it remains possible that cross-reactivity is due to a more complex mixture of mimetics, or very weak binding to empty HLA.

The importance of very high target affinity for this effect on therapeutic window was further supported by experiments with a TCR that binds its cognate MAGE-A3 pHLA with a weaker K_D_ of 17 nM and a t½ of 3.6 mins at 37°C (Table S2). This TCR had previously been used in a T cell therapy clinical trial that was discontinued due to unpredicted cross-reactivity to a Titin-derived mimetic.^23, 24^ Our measurements show this Titin mimetic bound the TCR with an affinity of 183 nM and a t½ of 0.46 mins at 37°C (Table S2). The fast off-rate of this TCR when binding its target suggests that dark-state formation is not likely to be a significant factor; and indeed, in contrast to TCR-X, a gain of potency was seen with the high affinity E24 variant on both target and mimetic without a large drop in efficacy, while the weaker binding E2 variant reduced potency with both the target MAGE-A3 and mimetic Titin peptides. This demonstrates that weakening CD3 binding only improves the therapeutic window when target pHLA binding has a very high affinity.

### Optimum CD3 affinity reflected in T cell cytotoxicity as well as IFN _γ_ release

The experiments in this manuscript have primarily focused on measuring IFNγ through direct measurement by ELISpot and ELISA. However, cytokine release is only one aspect of the T cell response and for CD8 T cells cytokine release has been shown to be much less sensitive to activation than cytotoxic responses such as microtubule reorganisation and release of perforin^25^. To investigate the effect of CD3 binding affinity on cytotoxicity, killing assays were carried out with both TCR-X and anti-NY-BR-1 ImmTAC molecules.

The due to the cross-reactivity profile of the TCR, experiments with TCR-X focused on testing intermediate and weak CD3 binding affinity variants in several different cytokine release assays (Figure S7 A and B) and an Incucyte killing assay (Figure S7 C). These results confirmed optimum activity was seen with the E28 variant, or kinetically similar E30 variant, with both cytokine release and killing.

A panel of anti-NY-BR-1 ImmTAC molecules with the full range of CD3 affinities was assayed against CAMA-1 A2B2M antigen-positive and SKMEL28 A2B2M antigen-negative cell lines in an Incucyte killing assay (Figure 7 A and S6 A). Again, E28 performed best, supporting a similar kinetic optimum for both killing and IFNγ release. In these assays though the high-affinity anti-CD3 variants E42, E24 and E17 all reached comparatively higher Emax levels, as measured by area under the curve analysis of killing up to 48 hrs. Killing assay protocols use significantly more effector cells than IFNγ ELISA and ELISpot assays to get a clear readout (10:1 instead of 1:1), but an ELISA carried out on media samples saved at the end of the killing experiment confirmed the discrepancy between killing and cytokine release with high affinity CD3 binding (Figure 7 B). Similar results were seen with a different batch of effector PBMC using an alternative Phenix killing assay that could monitor both living and dead cells (Figure 7C and S6 C).

**Figure 7.**
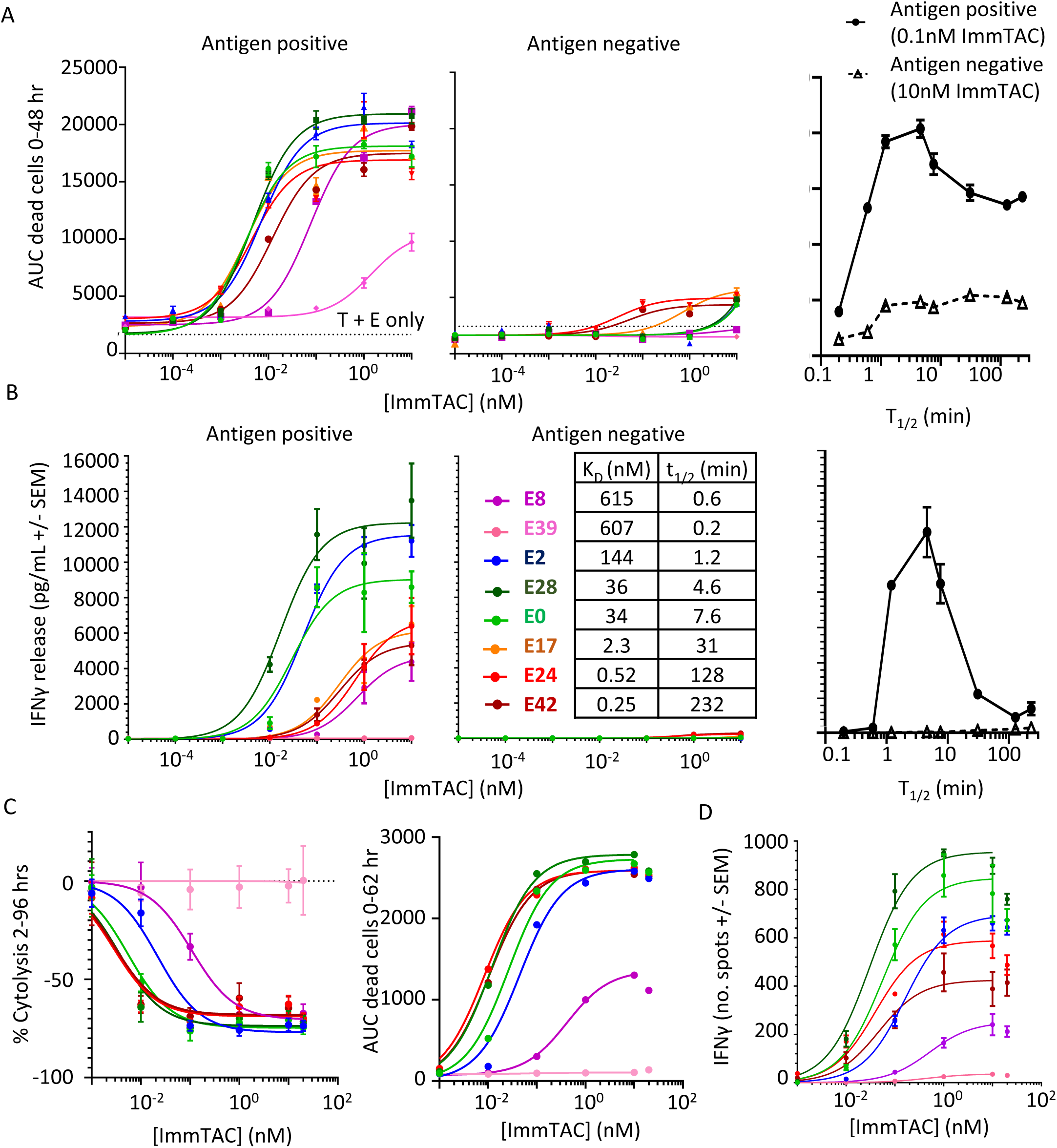
Comparison of cytotoxicity to cytokine release. **A)** Killing of antigen positive cells (left) and antigen negative cells (middle) with different CD3 affinity molecules as measured by an Incucyte assay. Area under the curve (AUC) of dead target cell counts at specified ImmTAC concentration. The plot on the right shows AUC vs t_1/2_ of CD3 binding at a singular ImmTAC concentration. Detailed time courses from which these data are derived are shown in Figure S6 **B)** Plot of IFNγ in media at the end of killing experiment measured by ELISA with antigen positive cells (left) and antigen negative cells (middle). **C)** Shows results from an alternative killing assay setup utilising Phenix instrument using CAMA1 antigen positive cells, but a different PBMC donor with a 4:1 ratio of effectors to targets. The left panel shows the percentage of surviving cells after 96 hrs relative to a control incubation without ImmTAC and normalised to initial cell counts at 2hrs (Detailed time courses shown in Figure S6), while the right panel uses data from counting dead cells and estimating the area under the curve in the first 62 hours of the experiment. **D)** ELISpot data collected with CAMA1 cells and the same PBMC donor as C, but with a 1:1 Effector to Target ratio.

While interesting these observations do not undermine the earlier conclusions that a receptor dark state supports an optimum dwell time for T cell triggering. If cytotoxicity is more sensitive to early triggering, and responds more rapidly to T cell stimulation, it might be less sensitive to dark-state formation than cytokine release. It should also be considered that the output of a cytotoxicity assay may be more limited in its dynamic range as there are only a certain number of target cells available to kill in each assay and they provide a more binary readout of T cell activity. Similarly the kinetics of the killing process is more limited by rate of T cell movement, as well as other factors, whereas cytokine release is ultimately limited by the translational output of the T cells, creating many potential discrepancies in the dynamic range of the response. Accurately modelling changes in pHLA presentation and other effects of cell killing would demand a significantly more complex model that fell outside the scope of this project. However, future studies might be designed to investigate these subtle distinctions between cytokine release and cytotoxicity, as well as conducting carefully designed *in vivo* studies to assess what combinations of these effects are most beneficial for reducing tumour burden.

## Discussion

To design an effective T cell engaging bispecific molecule, it must be made sufficiently potent to redirect patient T cells towards target cells at the chosen dose, while not triggering activation towards normal cells. A potentially dose-limiting toxicity observed with many T cell redirecting therapies is cytokine release syndrome.^26^ Stronger CD3 binding has previously been associated with increased incidence of systemic cytokine release and a CD38xCD3 bispecific antibody with unacceptable toxicity in non-human primates became better tolerated and more efficacious when the CD3 binding portion was engineered to weaken binding.^27^ Similarly, a CLL-1xCD3 bispecific with weak CD3 binding gave reduced systemic cytokine response compared to a strong CD3 binder,^28^ and both in vitro and in CD3-humanized mice, a HER2-targeted CD3 bispecific with strong CD3 binding induced greater cytokine production than one with weaker CD3 binding, despite comparable in vivo antitumor activity.^29^ One possible contributor to this effect could be the relationship we have demonstrated here between strong CD3 binding and increased responsiveness to weak off-target interactions.

Naturally occurring TCRs typically bind cognate HLA molecules with weak affinities, (micromolar range). However, the ImmTAC platform uses TCRs engineered for strong binding, (typically with picomolar affinities), while maximising the affinity window between target and mimetic pHLAs. The use of highly specific, picomolar-affinity TCRs here has enabled exploration of a range of binding strengths and specificities out of reach to previous studies. Molecules that target pHLA with intermediate affinity require stronger affinity CD3 binding to increase potency, which risks boosting cross-reactivity to weakly binding mimetic pHLAs that are often difficult to detect and characterise. The use of very strong affinity TCRs, on the other hand, allows tuning of anti-CD3 kinetics and opens up a valuable therapeutic niche; one where on-target potency and efficacy can be maximised, while minimising off-target activation by mimetic pHLAs. ImmTAC molecules already go through a rigorous preclinical safety package during their development, establishing the highly specific potential of this technology,^30^ and tuning anti-CD3 kinetics on a case by case basis makes it likely more therapeutic molecules will be able to meet these stringent criteria.

In this study, we have exclusively used bispecifics of the ImmTAC format, maintaining the same spatial relationship between the target-engaging and CD3-engaging moieties. Activity of CD3 bispecifics can be affected by factors such as distance of target epitope from the cell membrane,^12, 31^ CD3 epitope,^32^ and target distribution/mobility.^33^ As such, while the model is anticipated to be broadly applicable, the parameters identified by the model are likely to be specific to the conditions tested. For the vast array of other bispecific formats, with differing spatial arrangements of target and CD3 binding,^34, 35^ the optima for target and CD3 affinity may vary. Additionally many antibody based bispecific therapies bind to cancer surface antigens on target cells rather than pHLA which will alter their mode of action, as, unlike pHLA, these antigens are unable to recruit CD4 and CD8 co-receptors, and so may display significantly different optima.

We have shown that for both target engagement and CD3 engagement strong binding does not always improve potency and can be detrimental to T cell activity when the other arm of the bispecific also binds strongly. This is inconsistent with previous models of activity of CD3 bispecifics based on their binding kinetics,^36–39^ which have assumed a direct relationship between T cell activation and the concentration of the trimeric complex formed between CD3, bispecific, and target. In such a model, there is no optimal CD3 affinity for on-target activity, as stronger binding will always result in a higher concentration of trimer and greater T cell activation. This contrasts with conventional models of T cell activation where it is clear that the number of CD3 bridging interactions is not the sole determinant of activity,^15^ and the duration of bridging interactions plays a key role. The concept of HLA:TCR bridging interactions having an optimal dwell-time is well-established,^40^ with the upper limit variously explained by sustained signalling or limited signalling. Various mathematical models of T cell activation have been developed to explore the nature of this relationship.^14, 20, 41–43^ This concept has been invoked in the context of bispecific T cell engagers,^44^ but no corresponding mathematical model has previously been published.

The observation reported here that the optimum CD3 affinity for maximum response depends on target affinity, indicates a mechanism for improving specificity that is also incompatible with models driven purely by the concentration of target, biologic and effector. These experiments demonstrate that the affinity of CD3 engagement can be selected to maximally differentiate between high-affinity on-target interactions and low-affinity off-target interactions, to improve therapeutic window.

Our studies have focused on IFNγ release and cytotoxicity as the main outputs of T cell activation. It is possible that other cytokines could be affected differently. However, a recent study suggests that TNF-α and IL-2 release both share a similar activation threshold to IFNγ and respond similarly to co-stimulation.^45^ The differing sensitivity of T cell killing and cytokine release is well documented,^46^ and a number of studies have recently suggested the possibility of uncoupling cytotoxicity and cytokine release.^32, 37^ It could be hypothesised that if cytotoxicity is more sensitive to early triggering, and responds more rapidly to T cell stimulation, it might be less sensitive to dark-state formation than cytokine release. However, as the optimum anti-CD3 kinetics were the same for both cytokine release and cytotoxicity with two different TCRs, this suggests effective uncoupling cannot be achieved by altering affinity alone.

The affinity of a bispecific for CD3 has additional effects beyond those assessed here, including effects on the biodistribution and clearance of the bispecific molecule.^28, 47^ This manuscript focuses solely on the effects of affinity on T cell activation *in vitro* and when seeking to maximise potency and efficacy *in vivo* the role of biodistribution may well out-weigh concerns over optimising for T cell activation. Meaningful *in vivo* studies can be challenging to design for Immunotherapies, owing to the many sequence differences between human patients and model organisms. However, at this time only *in vivo* studies have the potential to comprehensively model clinical outcomes, and factor in the many different processes that may help and hinder ImmTAC activity in the body.

Understanding the trade-offs between the different effects of CD3 affinity through all means available should allow future development of safer, more effective therapeutics. With promising results with T cell redirecting molecules in clinical trials,^5, 48–51^ and many more in the pipeline, the age of bispecific therapies appears to be dawning. This study provides valuable information for developing the next generation of bispecific molecules, helping these game-changing drugs realise their maximum clinical potential.

## Materials and methods

### Proteins and affinities

ImmTAC fusion proteins were expressed in the BL21 (DE3) Rosetta pLysS strain, and refolded from inclusion bodies and purified as previously described.^52^ Purity was checked by reducing and non-reducing SDS-PAGE and concentrations assessed by A280 measurement and extinction coefficients derived from sequence by the inbuilt DNAdynamo algorithm. Peptide HLA molecules were also produced as previously described.^53^

Binding kinetics and affinities were measured by surface plasmon resonance by amine coupling CD3εδ expressed by HEK293 cells using a knob-in-holes Fc format (ACRO biosystems CDD-H52W0) to the CM5 sensor chip. Experiments were carried out using the single cycle protocol of the Biacore T200 system at 37 °C in PBS with 0.005% surfactant P20 with 5mM HCl used for chip regeneration between each experiment.

### Cell lines and cell culture

Cell lines used in this study were grown according to the manufacturers’ instructions. CAMA-1 cells (ATCC^®^ HTB-21^™^) express a low level of HLA-A2 that result in very low surface levels, thus this cell line was transduced in-house with a lentivirus containing HLA-A2 and beta 2-microglobulin (CAMA-1 A2b2m) for assays with ImmTAC-NY-BR-1. SKMEL28 A2B2M cell line was produced in the same way. Cell line authentication and mycoplasma testing were routinely carried out by the LGC Standards Cell line Authentication Service (www.lgcstandards.com) and Mycoplasma Experience Ltd (www.mycoplasma-exp.com), respectively. Peripheral blood mononuclear cells (PBMCs) were obtained from StemCell (lot SC003 1807130132, HLA-A0101,A0301) and Discovery Life Sciences (lot CB004Z1110036544120518A, HLA-A0301,A1101).

### IFN**γ** ELISPOT and ELISA

IFNγ ELISpot and ELISA assays were carried out according to the manufacturer’s instructions (BD BioSciences,Cat #552138; R&D, cat # DY285B; and Glo reagent,cat # DY993. Either in 384 well (12500 T2 and 12500 PBMC), or 96 well format (50,000 T2 and 50,000 PBMC). T2 cells were pulsed with peptides, either by adding NY-BR-1 specific peptide or irrelevant TAX peptide directly into the microplate wells at final concentrations ranging from 2.5 uM to 5 nM. For ELISpot plates were incubated overnight at 37 °C/5 % CO2 and quantified using an automated ELISpot reader (Immunospot Series 5 Analyzer, Cellular Technology Ltd.) or for ELISA plates were incubated for 48 hrs before developing using the R&D, cat # DY285B protocol and reading with an Enspire plate reader.

### Jurkat Luciferase assay

The same 96 well setup was used as for the PBMC assay with 50000 T2 and 50000 Jurkat NFAT-Luciferase cells per well (Promega) in 100ul R10 medium. Plates were incubated for 22 hours at 37 °C/5 % CO2 prior to addition of 33ul Bio-Glo reagent (Promega G7940), then luminescence was read on a Clariostar plate reader.

### Incucyte S3 killing assay

NY-BR-1 antigen positive cell line CAMA-1 A2B2M and antigen negative SKMEL28 A2B2M were stained with CellTracker™ Deep Red Dye (Invitrogen, C34565) and seeded at 15000 and 10000 cells/well, respectively, in a 96 well cell culture plate and left to rest for 24 hours. NY-BR-1 anti-CD3 variant ImmTAC molecules were titrated and added together with HLA-A*02:01-negative PBMC donor (CB004) at a 10:1 E:T ratio. Target cells and effectors were stained with a green Caspase 3/7 dye for tracking cell apoptosis (IncuCyte Caspase-3/7 Green Apoptosis Assay Reagent,Essen Bioscience, 4440). Cells were incubated for 96 hours in the Incucyte S3 Live-Cell Analysis System (Essen Bioscience, 4647), which uses real-time quantitative live cell imaging, based on fluorescent nuclear staining of the Caspase-3/7 activated dye, to track the numbers of apoptotic cells at regular time points. Plates were removed from the incubator after 96 hours and supernatant was collected for IFNγ ELISA. The results of the killing assay were analysed with the Incucyte software and Graph Pad Prism used to plot 48 hour Area Under the Curve of killing over different ImmTAC concentrations.

### Phenix Killing assay

The Opera Phenix system (Perkin Elmer) was also used to follow both target cell growth and killing simultaneously. In this assay an effector : target ratio of 4:1 was used. Target cells were counted and plated at 15,000 cells/well in 100 uL/well (CAMA1-A2B2M) on the day prior to setting up the assay PBMCs were thawed from liquid nitrogen on day 2 counted, stained with 2 uM CellTracker DeepRed and plated in 50uL/well. NucView488 reagent was added to effectors to give a 0.8 uM final concentration on the well. Hoechst was added to effectors to give a final concentration of 150 nM n the well. Plates were imaged with the 5X objective every 6h over 96h.

### Mathematical modelling

The mathematical model was built using Matlab and parameter inputs described in Table S3. Cell numbers, well area and values for CD3 and HLA copy numbers per cell were used to calculate effective local concentrations at the bottom of the experiment well for a section with a width of 1 micron and establish initial conditions for a system of 10 ordinary differential equations (ODEs), or 15 ODEs in the case of the cross-reactivity model (complete sets of ODE equations are included in supplementary material). The local concentration of ImmTAC was included in the differential equation system and was replaced from the bulk solution at a rapid rate which did not lead to any significant depletion of available ImmTAC in these models. For most fitting and modelling the ODE23s solver was used over a time course of 48 hrs for ELISA experiments and 24 hrs for ELISpot, except with the cross-reactive model where ODE15s was used.

To convert the ODE solutions into the quantity of IFNγ expected at the end of the experiment, the local concentration of signalling complexes was converted to a rate of IFNγ production expected at that moment by applying a hill function with a hill slope of 1 and fitting a maximum level of production (IFNmax) and a sensitivity level (IFNC50).

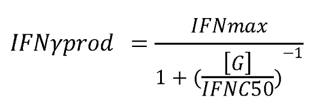

This equation was applied for the entire output vector and the area under the curve for the relevant time period was then calculated using the ‘trapz’ matlab function to obtain a total amount of IFNγ produced in that time. This method does not explicitly account for degradation or reuptake of IFNγ, and so any effects of these processes will be incorporated into the fitted IFNC50 and IFNmax parameters.

Fitting the model to the data was performed using the ABC SMC method previously described.^54, 55^ 1800 particles were generated at each iteration. Upper bounds and lower bounds were set depending on best guesses and are listed next to the fitted values in Supplementary Tables S4 and S5. Fit was assessed by combining the squared difference of simulated values from the mean of measured data plus standard deviation, meaning that any simulated values falling within the standard deviation of the data were considered to give a fit with no penalty. A minimum standard deviation of 10 pg/ml was applied to ELISA data and 5 spots to ELISpot data to account for any potential error not appropriately measured by the experiment.

## Acknowledgments

All work was funded by Immunocore LTD. The Authors would like to thank Rupert Kenefeck for many insightful discussions and guidance of cellular assays, Robert Pengally for his assistance refolding pHLA material, Laure Humbert for her assistance with preliminary cell assays, and Michelle McCully for their helpful comments and edits to this manuscript.

## Author Contributions

Conceptualization: I. B. R., R.M., N.D., A.V., R. O., S. H. and P.B.K.; Formal analysis: O.D. and I.B.R. Investigation: I.B.R., R.M., N.D., A.V., M.C., R.O. and F.A.; Writing – Original Draft: I.B.R and P.B.K.; Writing – Review & Editing: I.B.R, P.B.K., O.D., and D.C.; Supervision: R. O., S. H. and P.B.K.

