## Supplemental material for "Tuning the potency and therapeutic window of ImmTAC molecules by affinity modulation"

### Supplementary material

### Supplementary Tables

**Supplementary Table 1** – **Affinity of anti-CD3 for CD3εδ.** Biacore single-cycle measurements for anti-CD3 variants binding CD3εδ conducted at 37 °C in PBS supplemented with 0.005% surfactant P20.

| U number | $k_{on}$ | $k_{off}$ | CD3 binding $K_D$ (nM) | CD3 binding t 1/2 (min) |
|---|---|---|---|---|
| **E8** | 3.21E+04 | 0.0197 | **615** | **0.59** |
| **E39** | 1.18E+05 | 0.0714 | **607** | **0.16** |
| **E37** | 1.08E+05 | 0.0308 | **286** | **0.38** |
| **E41** | 9.38E+04 | 0.02472 | **264** | **0.47** |
| **E2** | 6.56E+04 | 0.009428 | **144** | **1.23** |
| **E35** | 1.02E+05 | 0.01396 | **137** | **0.83** |
| **E36** | 1.03E+05 | 0.01286 | **125** | **0.90** |
| **E31** | 9.01E+04 | 0.00811 | **90** | **1.42** |
| **E38** | 7.07E+04 | 0.00575 | **81** | **2.01** |
| **E30** | 7.39E+04 | 0.0028 | **37.8** | **4.13** |
| **E28** | 6.88E+04 | 0.00249 | **36.2** | **4.64** |
| **E0** | 4.48E+04 | 0.00152 | **33.9** | **7.60** |
| **E29** | 4.75E+04 | 0.00176 | **30.9** | **6.56** |
| **E16** | 1.37E+05 | 0.000812 | **5.91** | **14.23** |
| **E20** | 1.93E+05 | 0.00107 | **5.54** | **10.80** |
| **E17** | 1.60E+05 | 0.000373 | **2.34** | **30.97** |
| **E22** | 2.21E+05 | 0.000129 | **0.58** | **89.55** |
| **E24** | 1.74E+05 | 0.0000901 | **0.52** | **128.22** |
| **E42** | 2.03E+05 | 0.0000499 | **0.25** | **231.51** |

**Supplementary Table 2 – Affinity of TCRs used in this study for target and mimetic peptides.** Kinetics measured by Biacore at 37°C using single cycle method and five injections of tenfold dilutions. Target peptides presented by HLA are listed in peptide and sequence columns, including mimetic peptides with the amino acids that differ from the target highlighted in red. *MimH off-rate was potentially faster than the upper range that could be accurately measured by the instrument.

| TCR | Peptide | Peptide seq | HLA | $k_{on}$ | $k_{off}$ | $K_D$ nM | t 1/2 min | Notes |
|---|---|---|---|---|---|---|---|---|
| **NY-BR-1** | TAX | LLFGYPVYV | A2 | - | - | - | - | No binding |
| | MimI | SLIKNLDPV | A2 | - | - | 566900 | - | Equilibrium fit |
| | MimH | SLMKILSEV | A2 | *2.87E+5** | 1.89* | 65969 | *~0.006** | Equilibrium fit + dissociation used to calculate $k_{on}$ |
| | MimG | SLSIVLSTV | A2 | 3.85E+05 | 0.71963 | 1870 | 0.02 | |
| | MimF | SLSAILDTV | A2 | 3.69E+05 | 0.416805 | 1131 | 0.03 | |
| | MimE | SLSKITTTV | A2 | 5.44E+05 | 0.218521 | 402 | 0.05 | |
| | MimD | SLSKILATV | A2 | 7.33E+05 | 0.123204 | 168 | 0.09 | |
| | MimC | SLSKIADTV | A2 | 4.95E+05 | 0.043133 | 87 | 0.27 | |
| | MimB | SLSKALDTV | A2 | 3.10E+05 | 0.003494 | 12.3 | 3.3 | |
| | MimA | SLAKILDTV | A2 | 1.06E+06 | 0.001714 | 1.84 | 6.7 | |
| | **NY-BR-1** | **SLSKILDTV** | A2 | 3.81E+05 | 0.000156 | 0.37 | 74 | |
| **TCR-X** | | | A2 | 4.69E+05 | 0.000035 | 0.076 | 325 | |
| **MAG-IC3** | Titin | ESDPIVAQY | A1 | 1.37E+05 | 0.02493 | 183 | 0.46 | |
| | **MAGE-A3** | **EVDPIGHLY** | A1 | 1.92E+05 | 0.003193 | 16.7 | 3.6 | |

**Supplementary Table 3 – Parameters and model inputs**. Summary of variables input into the mathematical model, including which are fixed input values, and which are fitted to the data.

| Parameter | Symbol | Value |
|---|---|---|
| Well area | | 0.32 (96 wells) or 0.056 (384 well) |
| 'Active' volume | | 3.2000e-08 (96 well) Or 5.6000e-09 (384 well) (1 micron slice x area of well) |
| ImmTAC concentration | | 0.1 pM – 100 uM |
| Target cell number | | Number of target cells |
| T cell number | | Number of PBMC x 0.6 |
| CD3 per T cell | | 30,000 (Approximation base on literature and internaldata) |
| Target HLA per cell | | 332 for 5 nM pulse (Approximation based on internal epitope counting data) |
| Avogadro's number | | 6.0221409e+23 |
| ImmTAC Diffusion coefficient | D | 3.4e-8 (approximation that is too fast to allow meaningful depletion) |
| CD3 on rate | $k_{on}^{CD3}$ | See Table S1 |
| CD3 off rate | $k_{off}^{CD3}$ | See Table S1 |
| pHLA on rate | $k_{on}^{HLA}$ | See Table S2 |
| pHLA off rate | $k_{off}^{HLA}$ | See Table S2 |
| Time course | t | 48 hrs (ELISA)<br>24 hrs (ELISpot) |
| Membrane binding rate modifier | σ | fitted |
| Kinetic proof-reading rate | $k_p$ | fitted |
| Rate of dark state formation | $k_i$ | fitted |
| Rate of dark state recovery | $k_{rec}$ | fitted |
| Maximum IFNγ production | IFNmax | fitted |
| Sensitivity of IFNγ production | IFNC50 | fitted |

**Supplementary Table 4** – 'Best particle' fitted parameters from different NY-BR-1 experiments rounded to 3 significant figures. LB (Lower bounds) and UB (Upper bounds) of fitting are shown on the left.

| | | Units | LB | UB | Figure 1 | Figure 2 |
|---|---|---|---|---|---|---|
| Max iteration: | | | | | 25 | 20 |
| Membrane binding rate modifier | σ | none | $10^{-2}$ | $10^{4}$ | 0.294 | 0.074 |
| Kinetic proof-reading rate | $k_p$ | $\mu M^{-1} s^{-1}$ | $10^{-5}$ | $10^{2}$ | 0.0098 | 0.0023 |
| Rate of dark state formation | $k_i$ | $\mu M^{-1} s^{-1}$ | $10^{-4}$ | $10^{4}$ | 0.0102 | 0.097 |
| Rate of dark state recovery | $k_{rec}$ | $\mu M^{-1} s^{-1}$ | $10^{-15}$ | $10^{-1}$ | 0.00457 | 0.0000498 |
| Maximum IFN γ production | IFNmax | pg/ml $s^{-1}$ | $10^{-2}$ | $10^{4}$ | 39 | 124 |
| Sensitivity of IFNγ produ ction | IFNC50 | μM | $10^{-6}$ | $10^{1}$ | 1.23 | 0.97 |

**Supplementary Table 5** - 'Best particle' fitted parameters from PRAME experiments. For HLA2+ cells mimetic copies expected on both T cells and cancer cells and starting values for local concentration of off-target mimetic fitted as if displayed uniformly on both Target cells and T cells. LB (Lower bounds) and UB (Upper bounds) of fitting are shown on the left.

| | | Units | LB | UB | Donor 1 Figure 6 | Donor 2 Figure S4B | Donor 3 Figure S4C |
|---|---|---|---|---|---|---|---|
| | | | | | HLA-A201 + | HLA-A201 + | HLA-A201 - |
| Max iteration: | | | | | 25 | 20 | 23 |
| Membrane binding rate modifier | σ | none | $10^{-2}$ | $10^{2}$ | 4.87 | 0.439 | 0.0159 |
| Kinetic proof-reading rate | kp | $\mu M^{-1}\ s^{-1}$ | $10^{-5}$ | $10^{0}$ | 0.0013 | 0.00648 | 0.00143 |
| Rate of dark state formation | ki | $\mu M^{-1}\ s^{-1}$ | $10^{-5}$ | $10^{0}$ | 0.0144 | 0.00182 | 0.218 |
| Rate of dark state recovery | krec | $\mu M^{-1}\ s^{-1}$ | $10^{-8}$ | $10^{-3}$ | 2.67E-08 | 1.55E-06 | 5.33E-05 |
| Maximum IFNγ production | IFNmax | pg/ml $s^{-1}$ | $10^{-3}$ | $10^{4}$ | 0.0144 | 0.0247 | 14.9 |
| Sensitivity of IFNγ production | IFNC50 | μM | $10^{-7}$ | $10^{1}$ | 0.00000748 | 0.00129 | 0.00692 |
| Mimetic off-rate | | $s^{-1}$ | $10^{-4}$ | $10^{0}$ | 0.015 | 0.0108 | 0.319 |
| Mimetic copies per cell | | copies | $10^{1}$ | $10^{7}$ | 86.3 | 155 | 375 |
| Mimetic Affinity (μM) (kon fixed at 0.5 $\mu M^{-1}\ s^{-1}$) | | μM | | | 0.03 | 0.0216 | 0.638 |

Supplementary equations 1 – 10 ODE system:

**[A]** - Local ImmTAC in solution

$$\frac{d[A]}{dt} = -k_{on}^{CD3}[A][B] + k_{off}^{CD3}[C] - k_{on}^{HLA}[A][E] + k_{off}^{HLA}[D] - k_{off}^{CD3}[I] - k_{on}^{CD3}[A][J] - \frac{(D([A]-[Bulk]) \times area)}{activeVolume}$$

**[B]** - Free CD3

$$\frac{d[B]}{dt} = -k_{on}^{CD3}[A][B] + k_{off}^{CD3}[C] - \frac{k_{on}^{CD3}}{\sigma}[B][D] + k_{off}^{CD3}[F] + k_{off}^{CD3}[G] + k_{rec}[J]$$

**[C]** - CD3 bound to ImmTAC

$$\frac{d[C]}{dt} = k_{on}^{CD3}[A][B] - k_{off}^{CD3}[C] - \frac{k_{on}^{HLA}}{\sigma}[E][C] + k_{off}^{HLA}[F] + k_{off}^{HLA}[G] + k_{rec}[I]$$

**[D]** - Target HLA bound to ImmTAC

$$\frac{d[D]}{dt} = k_{on}^{HLA}[A][E] - k_{off}^{HLA}[D] - \frac{k_{on}^{CD3}}{\sigma}[B][D] + k_{off}^{CD3}[F] + k_{off}^{CD3}[G] + k_{off}^{CD3}[H] - \frac{k_{on}^{CD3}}{\sigma}[D][J]$$

**[E]** - Free target HLA

$$\frac{d[E]}{dt} = -k_{on}^{HLA}[A][E] + k_{off}^{HLA}[D] - \frac{k_{on}^{HLA}}{\sigma}[E][C] + k_{off}^{HLA}[F] + k_{off}^{HLA}[G] + k_{off}^{HLA}[H] - \frac{k_{on}^{HLA}}{\sigma}[E][I]$$

**[F]** - Fresh CD3-ImmTAC-target bridge

$$\frac{d[F]}{dt} = \frac{k_{on}^{HLA}}{\sigma}[E][C] - k_{off}^{HLA}[F] + \frac{k_{on}^{CD3}}{\sigma}[B][D] - k_{off}^{CD3}[F] - k_p[F]$$

**[G]** - Phosphorylated Bridges

$$\frac{d[G]}{dt} = k_p[F] - k_{off}^{HLA}[G] - k_{off}^{CD3}[G] - k_i[G]$$

**[H]** - Dark state target HLA bridge

$$\frac{d[H]}{dt} = k_i[G] - k_{off}^{HLA}[H] - k_{off}^{CD3}[H] + \frac{k_{on}^{HLA}}{\sigma}[E][I] + \frac{k_{on}^{CD3}}{\sigma}[J][D]$$

**[I]** - Dark state CD3 + IMMTAC-only

$$\frac{d[I]}{dt} = k_{off}^{HLA}[H] - k_{rec}[I] - k_{off}^{CD3}[I] + k_{on}^{CD3}[A][J] - \frac{k_{on}^{HLA}}{\sigma}[E][I]$$

**[J]** - Dark state CD3 only

$$\frac{d[J]}{dt} = k_{off}^{CD3}[I] - k_{rec}[J] - k_{on}^{CD3}[A][J] + k_{off}^{CD3}[H] - \frac{k_{on}^{CD3}}{\sigma}[J][D]$$

Supplementary equations 2 – 15 ODE system to define a system containing a cross-reactive mimetic:

**[A]** - Local ImmTAC in solution

$$\frac{d[A]}{dt} = -k_{on}^{CD3}[A][B] + k_{off}^{CD3}[C] - k_{on}^{HLA}[A][E] + k_{off}^{HLA}[D] - k_{off}^{CD3}[I] - k_{on}^{CD3}[A][J] - \frac{(D([A]-[Bulk]) \times area)}{activeVolume} - k_{on}^{mim}[A][K]$$

**[B]** - Free CD3

$$\frac{d[B]}{dt} = -k_{on}^{CD3}[A][B] + k_{off}^{CD3}[C] - \frac{k_{on}^{CD3}}{\sigma}[B][D] + k_{off}^{CD3}[F] + k_{off}^{CD3}[G] + k_{rec}[J] - \frac{k_{on}^{CD3}}{\sigma}[B][L] + k_{off}^{CD3}[M] + k_{off}^{CD3}[N]$$

**[C]** - CD3 bound to ImmTAC

$$\frac{d[C]}{dt} = k_{on}^{CD3}[A][B] - k_{off}^{CD3}[C] - \frac{k_{on}^{HLA}}{\sigma}[E][C] + k_{off}^{HLA}[F] + k_{off}^{HLA}[G] + k_{rec}[I] - \frac{k_{on}^{mim}}{\sigma}[K][C] + k_{off}^{mim}[M] + k_{off}^{mim}[N]$$

**[D]** - Target HLA bound to ImmTAC

$$\frac{d[D]}{dt} = k_{on}^{HLA}[A][E] - k_{off}^{HLA}[D] - \frac{k_{on}^{CD3}}{\sigma}[B][D] + k_{off}^{CD3}[F] + k_{off}^{CD3}[G] + k_{off}^{CD3}[H] - \frac{k_{on}^{CD3}}{\sigma}[D][J]$$

**[E]** - Free target HLA

$$\frac{d[E]}{dt} = -k_{on}^{HLA}[A][E] + k_{off}^{HLA}[D] - \frac{k_{on}^{HLA}}{\sigma}[E][C] + k_{off}^{HLA}[F] + k_{off}^{HLA}[G] + k_{off}^{HLA}[H] - \frac{k_{on}^{HLA}}{\sigma}[E][I]$$

**[F]** - Fresh CD3-ImmTAC-target bridge

$$\frac{d[F]}{dt} = \frac{k_{on}^{HLA}}{\sigma}[E][C] - k_{off}^{HLA}[F] + \frac{k_{on}^{CD3}}{\sigma}[B][D] - k_{off}^{CD3}[F] - k_p[F]$$

**[G]** - Phosphorylated target Bridges

$$\frac{d[G]}{dt} = k_p[F] - k_{off}^{HLA}[G] - k_{off}^{CD3}[G] - k_i[G]$$

**[H]** - Dark state target HLA bridge

$$\frac{d[H]}{dt} = k_i[G] - k_{off}^{HLA}[H] - k_{off}^{CD3}[H] + \frac{k_{on}^{HLA}}{\sigma}[E][I] + \frac{k_{on}^{CD3}}{\sigma}[J][D]$$

**[I]** - Dark state CD3 + IMMTAC-only

$$\frac{d[I]}{dt} = k_{off}^{HLA}[H] - k_{rec}[I] - k_{off}^{CD3}[I] + k_{on}^{CD3}[A][J] - \frac{k_{on}^{HLA}}{\sigma}[E][I] + k_{off}^{mim}[O] - \frac{k_{on}^{mim}}{\sigma}[K][I]$$

**[J]** - Dark state CD3 only

$$\frac{d[J]}{dt} = k_{off}^{CD3}[I] - k_{rec}[J] - k_{on}^{CD3}[A][J] + k_{off}^{CD3}[H] - \frac{k_{on}^{CD3}}{\sigma}[J][D] - \frac{k_{on}^{CD3}}{\sigma}[J][L] + k_{off}^{CD3}[O]$$

**[K]** - Free mimetic HLA

$$\frac{d[K]}{dt} = -k_{on}^{mim}[A][K] + k_{off}^{mim}[L] - \frac{k_{on}^{mim}}{\sigma}[K][C] + k_{off}^{mim}[M] + k_{off}^{mim}[N] + k_{off}^{mim}[O] - \frac{k_{on}^{mim}}{\sigma}[K][I]$$

**[L]** - Mimetic HLA bound to IMMTAC

$$\frac{d[L]}{dt} = k_{on}^{mim}[A][K] - k_{off}^{mim}[L] - \frac{k_{on}^{CD3}}{\sigma}[B][L] + k_{off}^{CD3}[M] + k_{off}^{CD3}[N] + k_{off}^{CD3}[O] - \frac{k_{on}^{CD3}}{\sigma}[L][J]$$

**[M]** - Mimetic HLA bound to IMMTAC bound to receptor

$$\frac{d[M]}{dt} = \frac{k_{on}^{mim}}{\sigma}[K][C] - k_{off}^{mim}[M] + \frac{k_{on}^{CD3}}{\sigma}[B][L] - k_{off}^{CD3}[M] - k_p[M]$$

**[N]** - Signalling mimetic bridge

$$\frac{d[N]}{dt} = k_p[M] - k_{off}^{mim}[N] - k_{off}^{CD3}[N] - k_i[N]$$

**[O]** - Dark state Bridge to mimetic HLA

$$\frac{d[O]}{dt} = k_i[N] - k_{off}^{mim}[O] - k_{off}^{CD3}[O] + \frac{k_{on}^{mim}}{\sigma}[K][I] + \frac{k_{on}^{CD3}}{\sigma}[J][L]$$

A

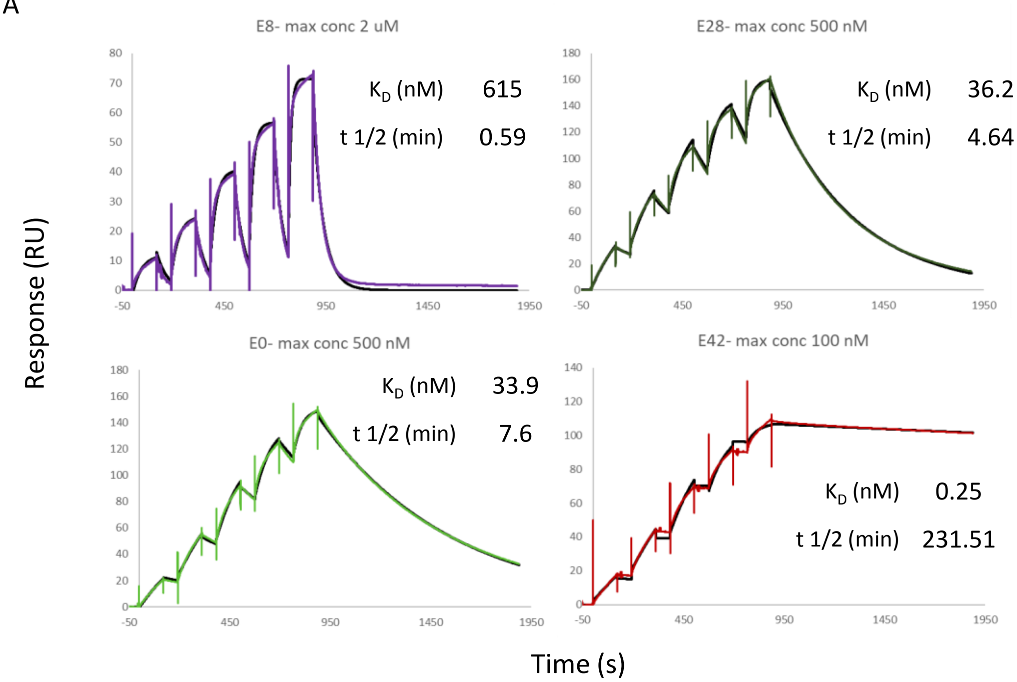


B

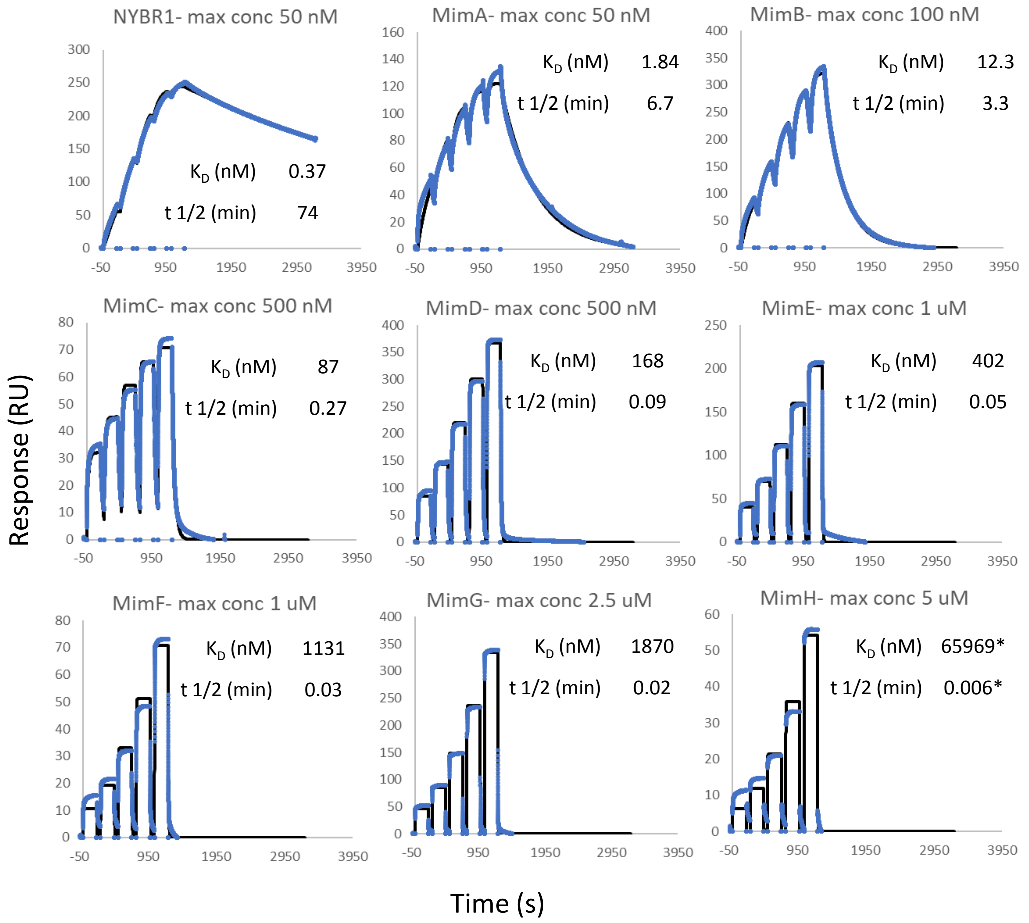


**Supplementary Figure S1 – Key SPR data. A)** Single cycle Biacore data for key anti-CD3 variants in the context of anti-NYBR1 ImmTAC molecules binding to human CD3εδ (ACRO biosystems) amine coupled to CM5 sensor chip at 37 °C in PBS pH 7.4 with 0.005% P20. Five 2x serial dilutions were injected with the highest concentration listed above each graph. Data is coloured according to variants and kinetic fits are shown with a black line. Data was collected with a Biacore T200 instrument. **B)** Single cycle Biacore data for anti-NYBR1 TCR binding to different biotinylated pHLA at 37 °C in PBS pH 7.4 with 0.005% P20. pHLA were immobilised via streptavidin that was amine coupled to a CM5 sensor chip and analyses carried out using a Biacore 8K instrument. *KD value determined by equilibrium fit and t1/2 based on separate fitting of off-rate, likely inaccurate due to the rapid dissociation observed.

A

| | $K_D$ (nM) | $t_{1/2}$ (min) |
|---|---|---|
| E8 | 615 | 0.6 |
| E39 | 607 | 0.2 |
| E2 | 144 | 1.2 |
| E31 | 90 | 1.4 |
| E38 | 81 | 2.0 |
| E28 | 36 | 4.6 |
| E0 | 34 | 7.6 |
| E16 | 5.9 | 14.2 |
| E17 | 2.3 | 31 |
| E24 | 0.52 | 128 |
| E42 | 0.25 | 232 |

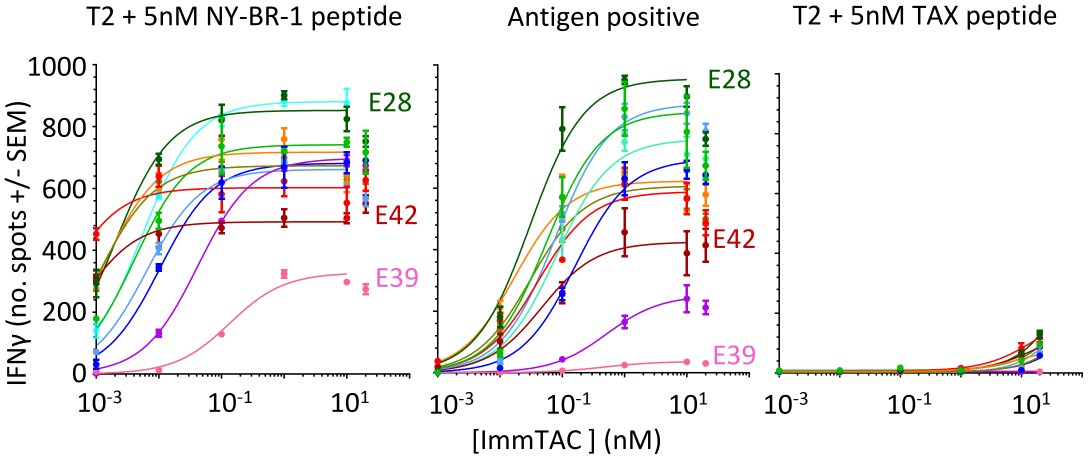


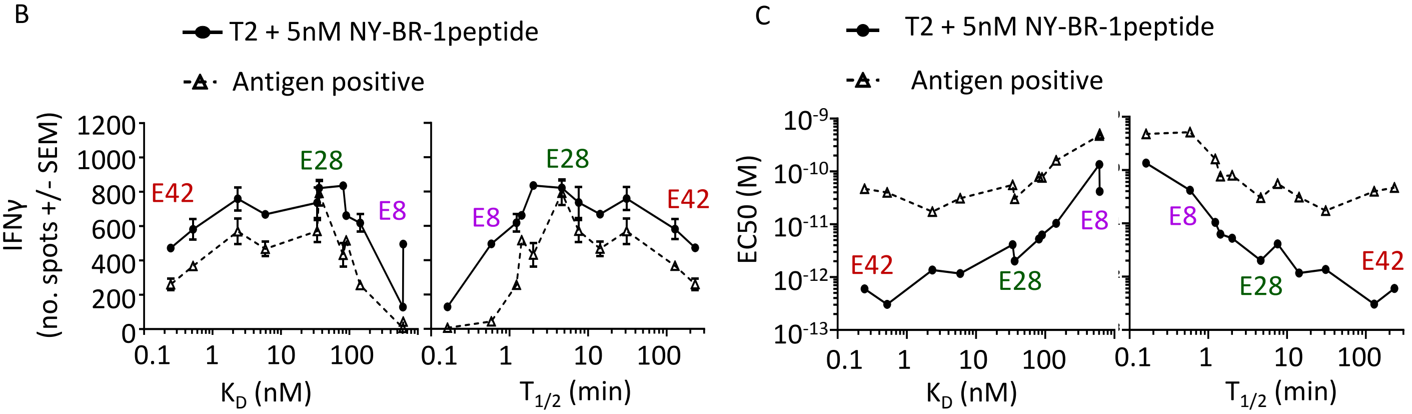


D

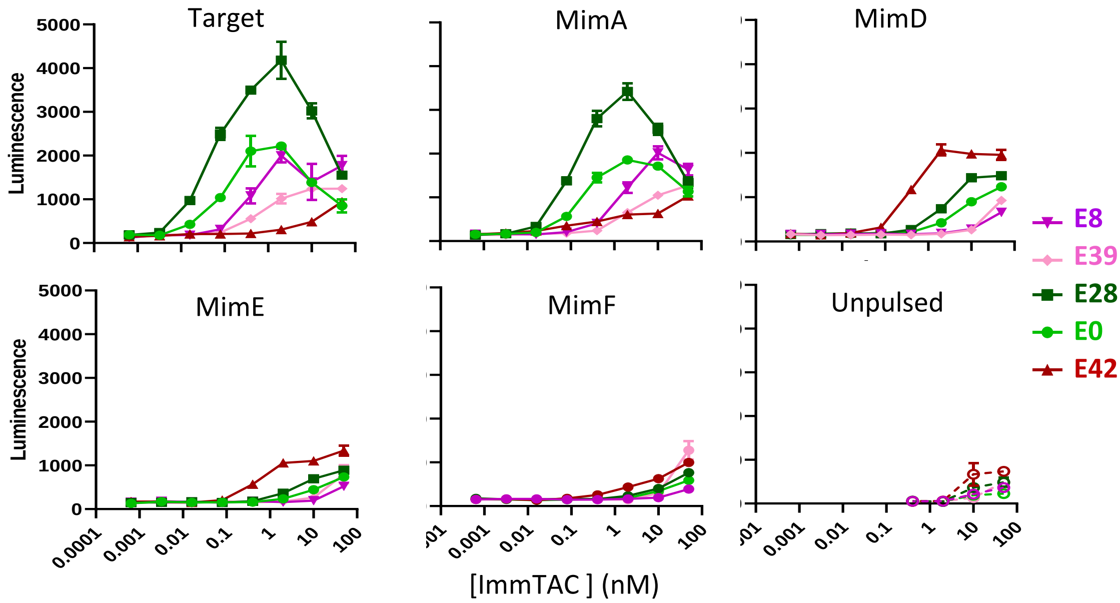


| | $K_D$ (nM) | $t_{1/2}$ (min) |
|---|---|---|
| E8 | 615 | 0.6 |
| E39 | 607 | 0.2 |
| E28 | 36 | 4.6 |
| E0 | 34 | 7.6 |
| E42 | 0.25 | 232 |

**Supplementary Figure S2 – Effect of CD3 affinity on ImmTAC activity with a highly specific TCR (NY-BR-1)**
**A)** IFNγ production measured by ELISpot from HLA-A*02:01 negative PBMCs in the presence of ImmTAC molecules with different anti-CD3 affinities from, left: T2 cells pulsed with 5 nM NY-BR-1 peptide, centre: antigen positive CAMA-1 A2B2M cells, or right: T2 cells pulsed with irrelevant TAX peptide. All data points representative of 3 experimental repeats. **B)** Summary plots of ELISpot data showing spots at 0.1 nM ImmTAC plotted against CD3 binding affinity or $t_{1/2}$. **C)** Summary plots of $EC_{50}$ plotted against CD3 binding affinity or $t_{1/2}$ . **D)** Activation assays with jurkat NFAT-luciferase reporter cell line (Promega) incubated for 22 hours with T2 cells pulsed with 1 nM target or mimetic peptides.

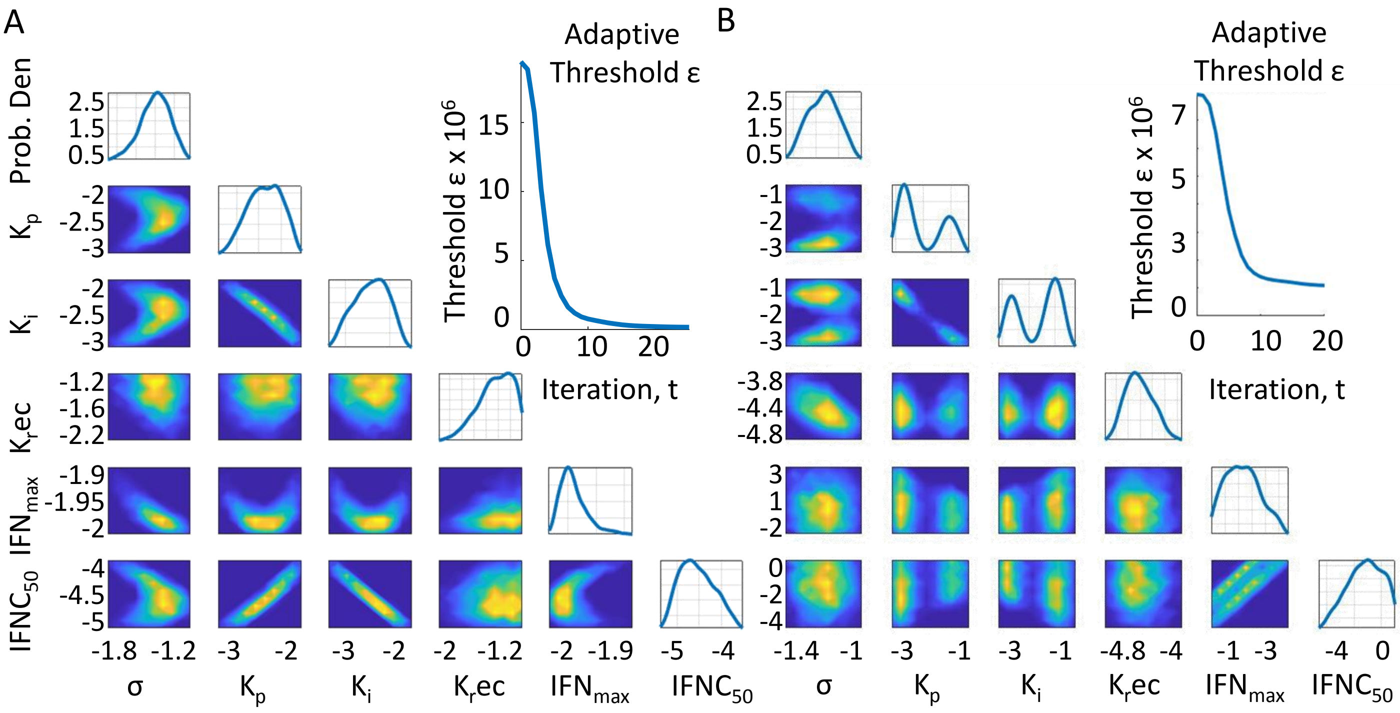


**Supplementary Figure S3 – Probability distributions for Model fitting. A)** Probability distributions for model parameters at final iteration of ABC-SMC fit shown in figure 4A, with values expressed as log values, ie 10 ^ (value on axis) and best particles values listed in table S1. **B)** Parameters distributions for fit to data shown in Figure 4B. A single optimum parameter set was not obtained here but two general solutions seem possible to fit the data. One with a faster kinetic proof-reading rate and a slower rate of dark state formation and the other with a fast dark-state formation rate and slower kinetic proof-reading rate. The fits to the data obtained with either parameter set both appear visually very similar and the same outlier data points are observed (Not shown).

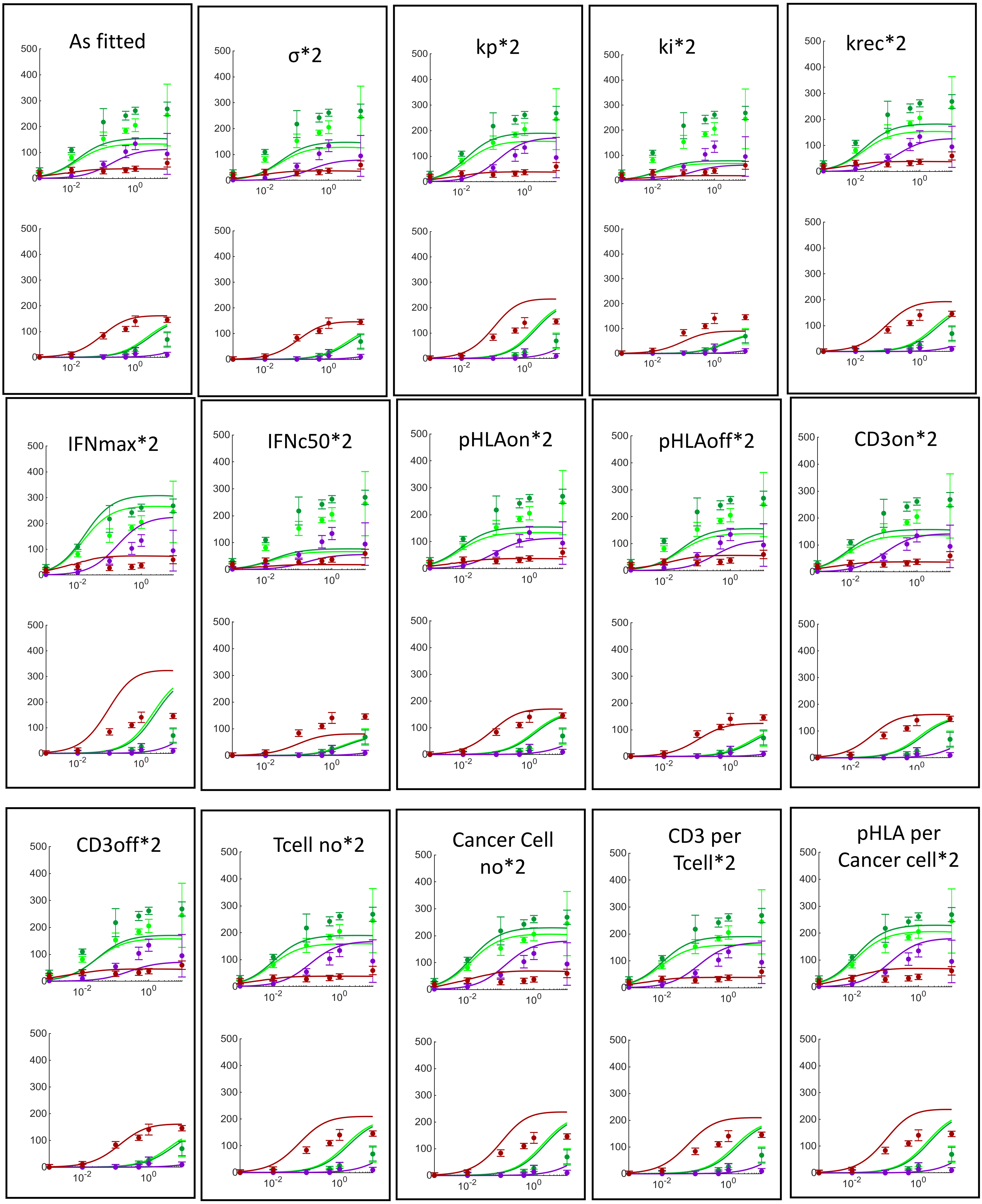


**Supplementary Figure S4– Effect of 2x change in fitted and fixed parameters on model output for NYBR1 Target and MimC.** To assess the sensitivity of the model to fitted and non-fitted parameters simulations were performed doubling the value of one parameter at a time and results are shown for target (top) and MimC (bottom). Changing some parameters appears to improve the fit with NYBR1-target reactivity but gives a worse fit with MimC. Generally however parameter changes on this scale do not significantly alter the output of the model and the trends remain the same.

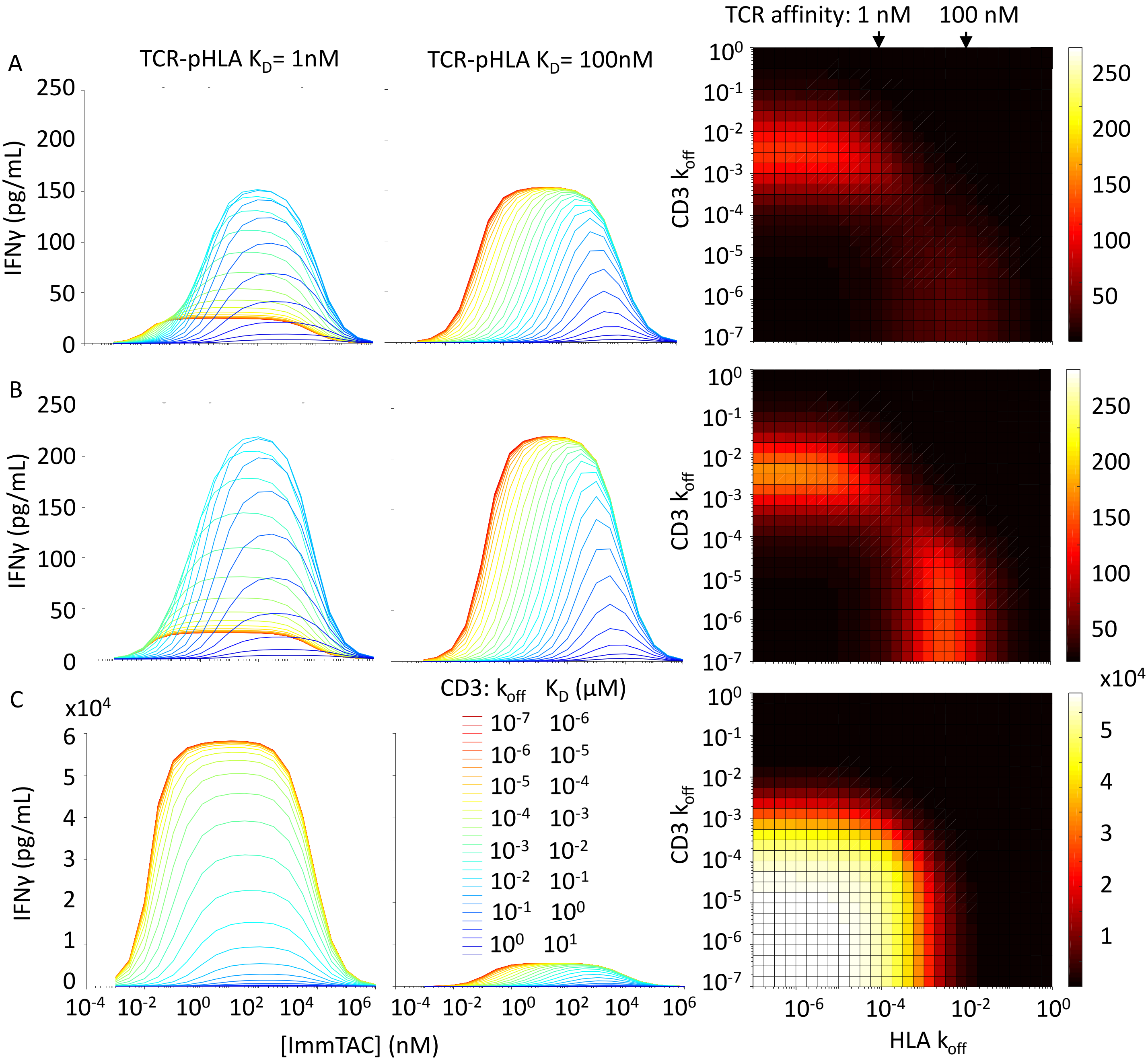


**Supplementary Figure S5– Effect of altering key model parameters. A)** Result of simulation using parameter fits from Figure 4B (Table S4) showing the effects of varying CD3 off-rate. Left: strong TCR-pHLA binding ($K_D$= 1nM), middle: intermediate TCR-pHLA binding ($K_D$= 100nM), and right: a heat map of response at 10 pM ImmTAC with different TCR-pHLA or CD3 affinity combinations. **B)** Effect of using same parameters as in A, but making dark-state recovery extremely fast when released from a complex ($k_{rec}$ = $10^{15}$). While overall response only increases slightly the main difference is that the response with intermediate TCR-pHLA : strong CD3 binding at 10 pM is more similar to that seen with strong TCR-pHLA : intermediate CD3 binding. **C)** Effect of using same parameters as in A, but making dark-state formation extremely slow ($k_i$= $10^{-30}$). Without dark-state formation a considerably higher response is generated with both strong and intermediate affinity TCR-pHLA binding, and strong TCR-pHLA : strong CD3 binding generates the highest responses. CD3 off-rates in all figures here range from $10^{-7}$ $s^{-1}$ (red) to $10^{0}$ (blue) with a constant on-rate of 0.1 $\mu M^{-1}$ $s^{-1}$.

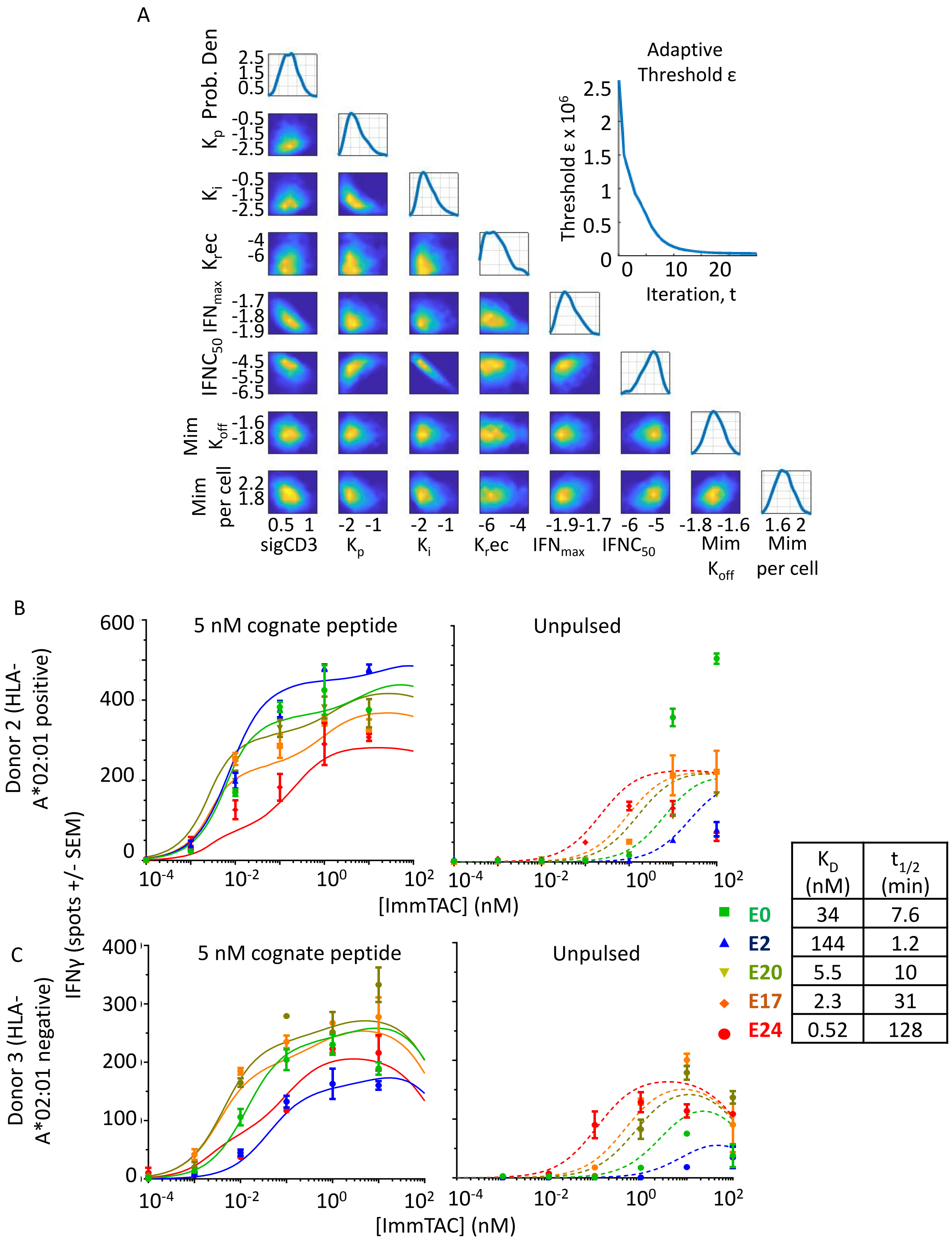


| | $K_D$ (nM) | $t_{1/2}$ (min) |
|---|---|---|
| E0 | 34 | 7.6 |
| E2 | 144 | 1.2 |
| E20 | 5.5 | 10 |
| E17 | 2.3 | 31 |
| E24 | 0.52 | 128 |

**Supplementary Figure S6– ABC-SMC fitting data for cross-reactivity model applied to tool ImmTAC with 'TCR-X'. A)** Parameter distributions obtained when fitting ELISpot data in Figure 6 B. Parameters expressed as log values, (ie 10^ value on axis) and best particle values are listed in Table S5. **B)** ELISpot of IFNγ production from a different PBMC donor (donor 2). Lines show model fits with parameters obtained in Table S5. **C)** ELISpot of IFNγ production from a third, HLA A*02:01 negative, PBMC donor (donor 3). Lines show model fits with parameters obtained in Table S5.

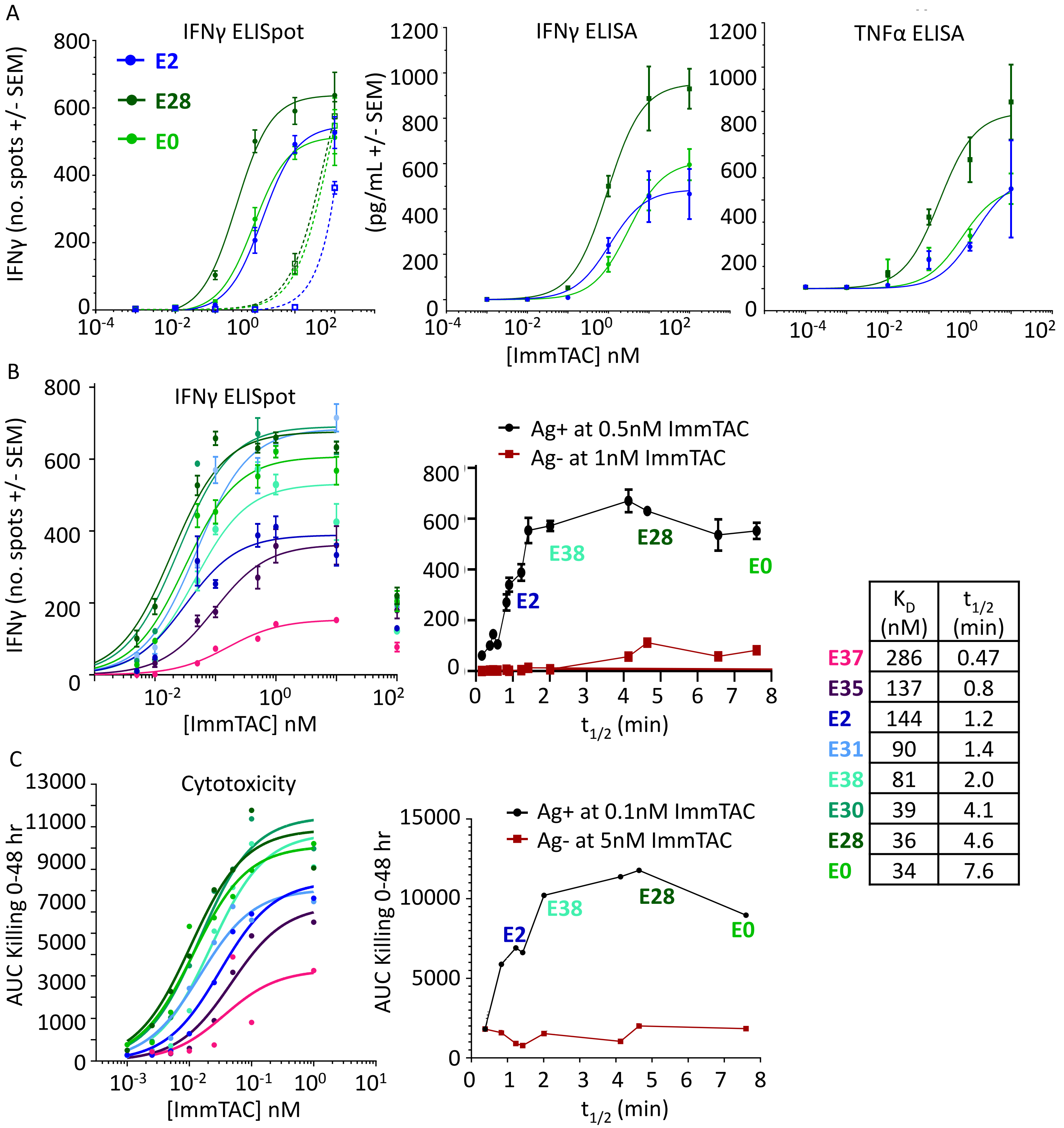


| | $K_D$ (nM) | $t_{1/2}$ (min) |
|---|---|---|
| E37 | 286 | 0.47 |
| E35 | 137 | 0.8 |
| E2 | 144 | 1.2 |
| E31 | 90 | 1.4 |
| E38 | 81 | 2.0 |
| E30 | 39 | 4.1 |
| E28 | 36 | 4.6 |
| E0 | 34 | 7.6 |

**Supplementary Figure S7 – Confirming optimal anti-CD3 kinetics on tool ‘TCR-X’ in cytokine release and cytotoxicity assays. A)** Focused comparison of E0, E28 and E2 variants. PBMCs (HLA-A*02:01 positive) incubated with antigen positive Mel624 cells (filled circles with solid lines) or antigen negative Granta519 cells (open squares with dashed lines). Curves represent 3-parameter fits to data obtained by IFNγ ELISpot (left), IFNγ MSD ELISA (middle), and TNFα MSD ELISA. **B)** Testing a focused panel of nanomolar affinity anti-CD3 variants: ELISpot on Antigen positive MEL624 cell line, with each data point performed in triplicate. The E28 variant was included on every plate and all data was normalised to the result obtained with this variant. Curves represent simple 3-parameter fits of data up to 10 nM ImmTAC. On the right the responses at specific ImmTAC concentrations are plotted against $T_{1/2}$ and also includes response to antigen negative MDAMB231 cell line. **C)** Incucyte killing assays of the same focused panel with areas under the curves up to 48 hours plotted against ImmTAC concentration. On the right AUC at a specified ImmTAC concentration is plotted against $T_{1/2}$ of CD3 binding, including response to antigen negative MDAMB231 cell line.

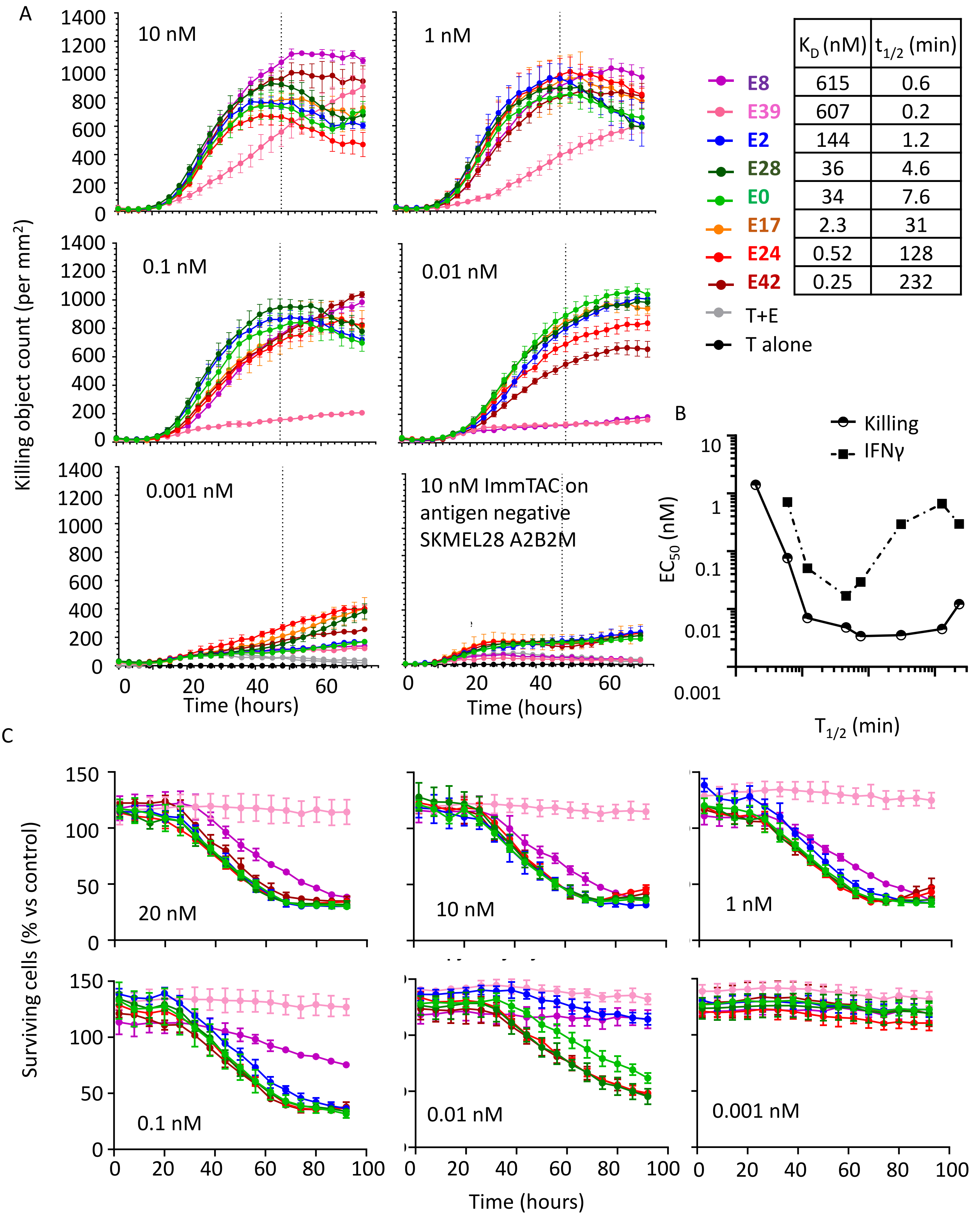


**Supplementary Figure S8 – Effect of CD3 affinity on killing of NYBR1 positive cells. A)** Killing time courses as measured by Incucyte assay with 8 different anti-CD3 variants. 5 panels show results on CAMA-1 A2B2M antigen positive cells with ImmTAC concentration shown top left, and one panel (bottom right) shows killing with the SKMEL28 A2B2M antigen negative cell line at 10 nM ImmTAC. **B)** Plot of $EC_{50}$ for IFNγ release against $t_{1/2}$ of CD3 binding. **C)** Data from a separate killing experiment using the Phenix instrument where here the % of surviving target cells is plotted relative to control target growth without ImmTAC.
